# Walking *Drosophila* aim to maintain a neural heading estimate at an internal goal angle

**DOI:** 10.1101/315796

**Authors:** Jonathan Green, Vikram Vijayan, Peter Mussells Pires, Atsuko Adachi, Gaby Maimon

## Abstract

While navigating their environment, many animals track their angular heading via the activity of heading-sensitive neurons. How internal heading estimates are used to guide navigational behavior, however, remains largely unclear in any species. We found that normal synaptic output from heading neurons in *Drosophila* is required for flies to stably maintain their trajectory along an arbitrary direction while navigating a simple virtual environment. We further found that if the heading estimate carried by these neurons is experimentally redirected by focal stimulation, the fly typically turns so as to rotate this internal heading estimate back towards the initial angle, while also slowing down until this correction has been made. These experiments argue that flies compare an internal heading estimate with an internal goal angle to guide navigational decisions, highlighting an important computation underlying how a spatial variable in the brain is translated into navigational action.

**One Sentence Summary:** Flies compare an internal heading estimate with an internal goal angle to guide navigation.

---

Many animals, from insects to mammals, keep track of their two-dimensional position (1, 2) and angular heading (*3, 4*) as they navigate through an environment. Neurons that track heading were first discovered in rodents (5), and more recently in insects (6, 7), including *Drosophila* (8). Whereas emphasis has been placed on understanding how the physiological properties of heading neurons are built (8-14), recent experiments have also begun to explore how animals use internal heading signals to guide navigational behavior (15). For example, destabilizing a rat’s head-direction system induces longer, more circuitous routes to a home position (16), suggesting that head direction cell activity is generally important for oriented navigation. Furthermore, electrophysiological recordings in flying bats have revealed not only neurons that track the bat’s head direction, but also neurons that track its *goal* direction––i.e., the angle of a known landing platform relative to the bat (17)––suggesting that a neural comparison between heading- and goal-direction guides the bat’s navigational behavior. However, whether such a neural comparison takes place and how the output of any such comparison is translated into navigational action remains poorly understood in any species. Here, we describe a behavioral task in which *Drosophila* maintain a consistent walking direction for minutes in a simple virtual environment. We further provide correlational and perturbational evidence that flies accomplish this task by turning so as to maintain a neural heading estimate at an internal goal angle, which can change direction over time.

To study how *Drosophila*’s heading system guides navigational behavior, we first developed a simple task that is likely to require a sense of direction. We placed tethered flies walking on an air-supported ball (9,18, 19) at the center of a cylindrical, 270° LED arena (20). We presented the flies with a bright, vertical bar whose position on the LED display rotated in closed-loop with the fly’s yaw (left/right) rotations on the ball, simulating a distant, static landmark (9, 21) (Figure 1A). In this virtual environment, we found that, without training, flies tended to walk forward while maintaining their heading relative to the vertical bar for several meters (Figure 1B, left). The flies sometimes also changed heading either abruptly or gradually over time (Figure 1B, center and right). Although previous studies have noted that *Drosophila* tend to fixate vertical stripes directly in front (22-25), the *arbitrary-angle fixation* we observed in our setup better resembles previous reports of flies maintaining an arbitrary heading relative to a visual object (*menotaxis*) (26) or to the angle of polarized light (27, 28), and is reminiscent of dispersal behaviors observed in the wild (29, 30). We have yet to fully characterize the specific experimental conditions that promote front-fixation vs. arbitrary-angle fixation in walking flies (see *Behavioral conditions* in Methods), but all genotypes we studied reliably performed arbitrary-angle fixation in our setups, allowing us to examine how this behavior is implemented at the neural level.

**Figure 1.**
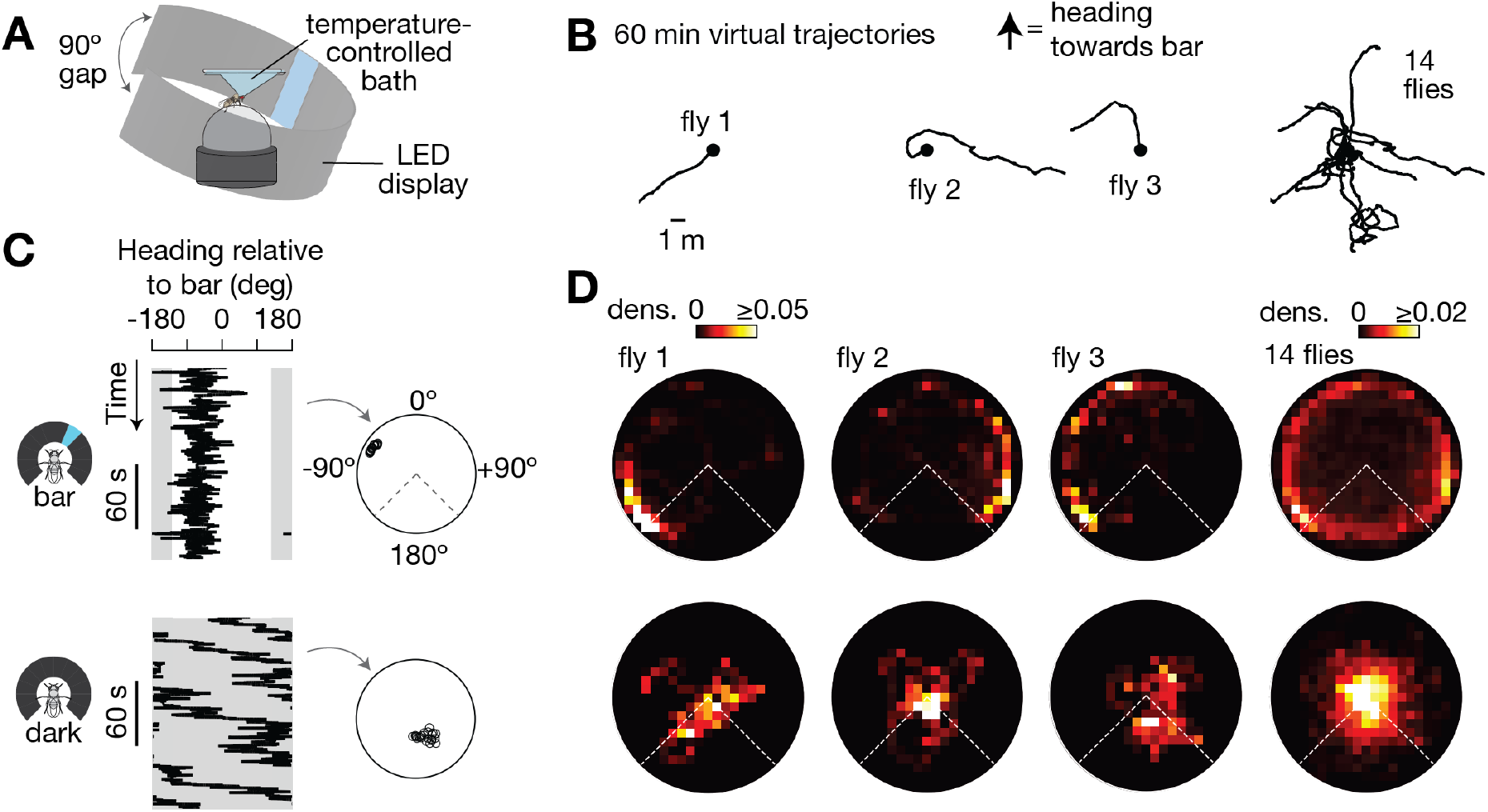
Walking flies maintain their heading at an arbitrary angle relative to a visual landmark for minutes. (**A**) Tethered fly walking on a ball at the center of a 270° LED arena. (**B**) 2D virtual trajectories of Canton-S wildtype flies walking with a bright bar in closed-loop. (**C**) Left: Heading relative to bar over 3 min. Right: We slid a 60 s analysis window over each heading trace and calculated the mean heading vector in each window (black circles). (**D**) Polar distributions of mean heading vectors taken over 60 s windows (slid by 1 s increments) for the flies in panel (B), above. Time points in which flies were standing still (i.e. forward velocity < 0.5 mm/s) were ignored for the calculation of each mean heading vector because heading values during such time points are stable for the trivial reason that the fly is not moving. Grey areas in (C) and dashed lines in (D) highlight the 90° gap of the LED display in which the bar is not visible.

To quantify the flies’ headings over time, we treated each heading measurement in a walking fly as a unit vector, computed the mean of unit vectors over 60 s windows (Figure 1C), and visualized the distribution of 60-s mean vectors in a polar plot (Figure 1D, see Methods). As expected from their virtual 2D trajectories (Figure 1B), individual flies tended to walk in a relatively constant direction with respect to the bar over 60 s (data points near the edge of the unit circle, Figure 1C-D, top). On the other hand, the same flies did not walk as straight in the dark (data points near the center of the circle, Figure 1C-D, bottom), even though their first-order walking statistics were similar in the dark and with the bar (Figure S1A), indicating that visual feedback allows the flies to stably maintain their direction. Beyond 60 s, flies maintained a consistent heading relative to a bar in closed-loop for many minutes (Figure S1B, top), but not in the dark (Figure S1B, bottom).

To determine whether flies actively maintain their heading in this simple virtual environment––rather than just walking straight irrespective of the location of the bar––we measured the behavioral responses of flies to experimentally introduced rotations of the visual scene. As the flies walked in closed-loop with a bar, we discontinuously rotated (or “jumped”) the bar ±90° or 180° relative to its current angular position. We operationally defined the fly’s *goal heading* as its mean heading over 10 s before bar jumps, analyzing trials where the flies’ headings were relatively constant (circular s.d. < 45°) before this experimental perturbation. We found that, following bar jumps, flies typically returned the bar to their goal heading (Figure 2A-B, Figure S2A). Moreover, flies performed similar corrections after 30 s of darkness (Figure 2C), indicating that they remembered their goal heading direction, for at least 30 s. That flies return the bar to a goal angle after 180° rotations (Figure S2A) and after 30 s of darkness, argues against the idea that they were responding solely to the angular velocity of the visual scene (i.e. stabilizing the bar’s yaw movements independent of its position on the arena). Rather, these behavioral data argue that flies actively maintain their heading with respect to a goal heading angle, whose angular position appears to be largely arbitrary, with a slight bias toward the rear-left and rear-right of the LED display (Figure S2B).

**Figure 2.**
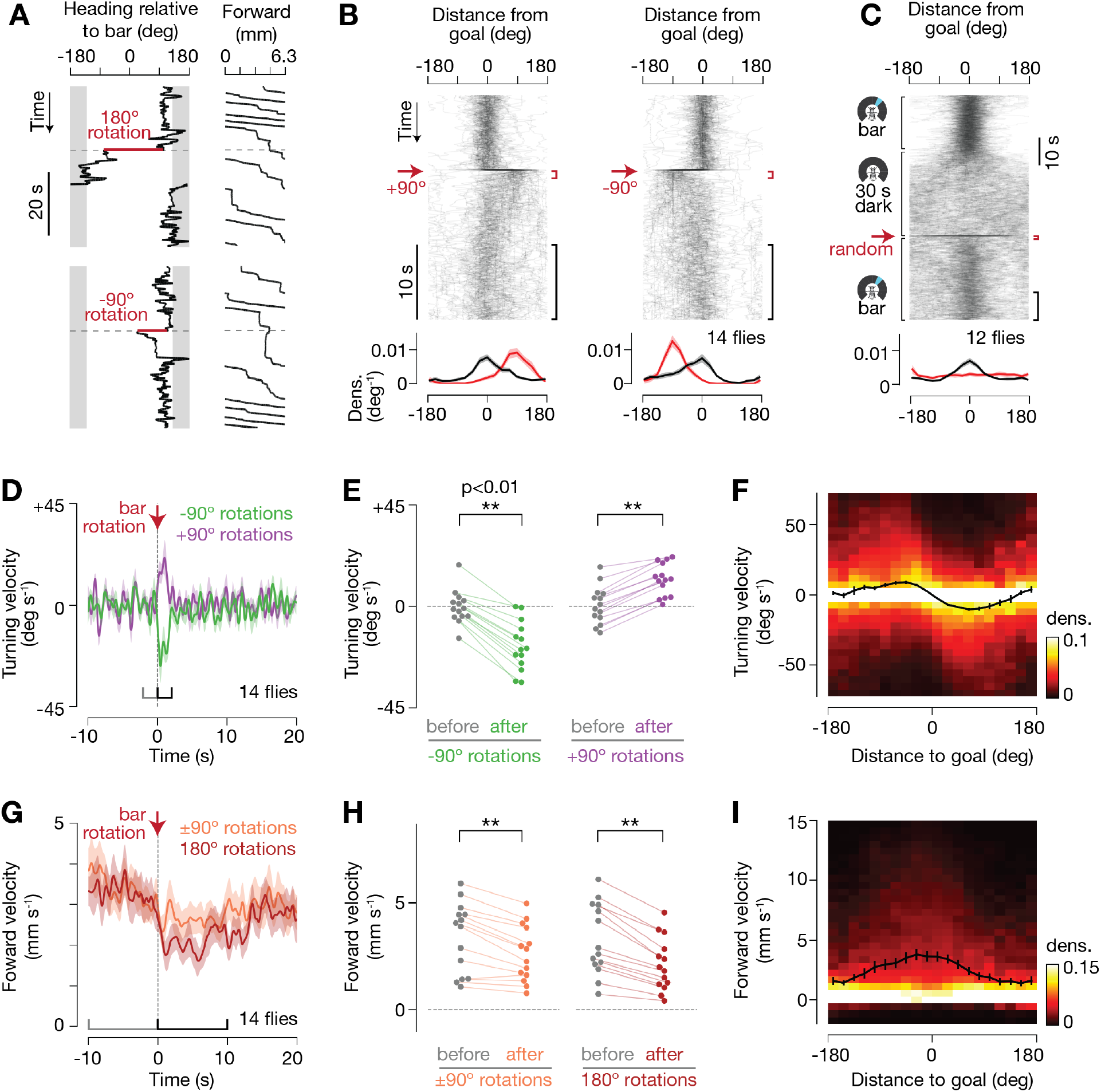
Flies actively turn towards an angular goal heading and resume fast forward walking once aligned with it. (**A**) Heading relative to bar and forward walking during bar jumps. 6.3 mm of forward walking is equal to one rotation of the ball (360°). (**B**) Top: Distance from goal over time for 90° bar jumps. The goal was operationally defined as the mean heading averaged across the 10 s window immediately prior the bar jump. In each panel, 90 traces from 14 flies are shown with 5% opacity. Bottom: Distributions of distance from goal at 0-1 s (red), and 10-20 s (black) after the bar jump. Mean and s.e.m. across flies are shown. (**C**) Top: Distance to goal over time for flies presented with a dark screen for 30 s, after which the bar reappeared at a random offset with respect to the ball. 167 traces from 12 flies are shown with 3% opacity. Bottom: Distance to goal distributions over 0-1 s (red) and 20-30 s (black) after the bar reappeared. (**D**) Turning velocity over time during 90° bar jumps. Mean and s.e.m. across flies are shown. (**E**) Mean turning velocity for each fly before (–2 to 0 s) and after (0 to 2 s) bar jumps. p-values were computed using the Wilcoxon signed-rank test. (**F**) Turning velocity as a function of distance to goal. Data from −20 s to 40 s around bar jumps were used to create this plot. Each column of the heat map is normalized independently because we had many more data points near x=0. Mean and s.e.m. across flies are shown (black curve). (**G-I**) Same as (D-F), but for forward velocity during 90° and 180° bar jumps. In (H), forward velocities were compared across 10 s windows immediately before and after bar jumps. For (B-I), we only analyzed trials where the flies maintained a relatively stable heading (circular s.d. < 45°) for 10 s before the bar jump, as an indication that flies were performing arbitrary-angle fixation. Approximately 10% of trials were excluded. No forward walking requirements were applied in this figure.

If flies navigate with respect to a goal direction, one might expect that both their turning and forward-walking velocities would vary systematically as a function of the difference between their current- and goal-headings. After experimentally induced ±90° rotations of the visual scene, flies tended to turn left or right so as to bring the bar back to its previous position on the arena (Figure 2D-E). Moreover, we observed a quantitatively varying relationship between turning velocity and the difference between the fly’s current- and goal-headings, with the largest mean, directional, responses evident when the bar was ±30-60° off the goal (Figure 2F).

Interestingly, we also observed that flies reduced their forward walking velocity after rotations in the visual scene (Figure 2G-H). Like with turning velocities, there was a systematic relationship between the flies’ forward-walking velocities and the angular difference between their current- and goal-headings (Figure 2I). Specifically, flies walked forward fastest when aligned with the goal, consistent with the flies attempting to effectively travel along their goal heading while reducing displacements away from it. The fact that flies slowed down after 180° rotations even when they turned very little (and therefore returned the bar very slowly) (Figure S3) argues that slowing down reflects a separable behavioral response from turning. Thus, two motor actions appear quantitatively tied to a comparison of the fly’s current- and goal-heading during navigation.

How might navigational signals in the brain allow flies to perform arbitrary-angle fixation? Recent work has identified several neuron classes in the fly central complex (Figure 3A-B) that carry heading signals. For example, E-PG (<u>e</u>llipsoid body-<u>p</u>rotocerebral bridge-<u>g</u>all) neurons (Figure 3B) (31) show a single calcium activity peak that rotates around the donut-shaped ellipsoid body (Figure 3C) (8), and 2-3 peaks that move left and right across the linear protocerebral bridge (9, 10), with the position of these peaks correlating with the fly’s virtual heading. To date, studies have focused on understanding how E-PG heading signals are updated by sensory and motor-related signals (8-10), but how E-PG activity guides navigational behavior remains unknown (Figure 3D). Given the behavioral evidence presented above, one possibility is that the E-PG heading signal is compared with an internal goal heading signal, and the angular difference, or error, between these two signals drives the fly to turn toward the goal direction and walk faster when aligned with it (Figure 3D-E).

**Figure 3.**
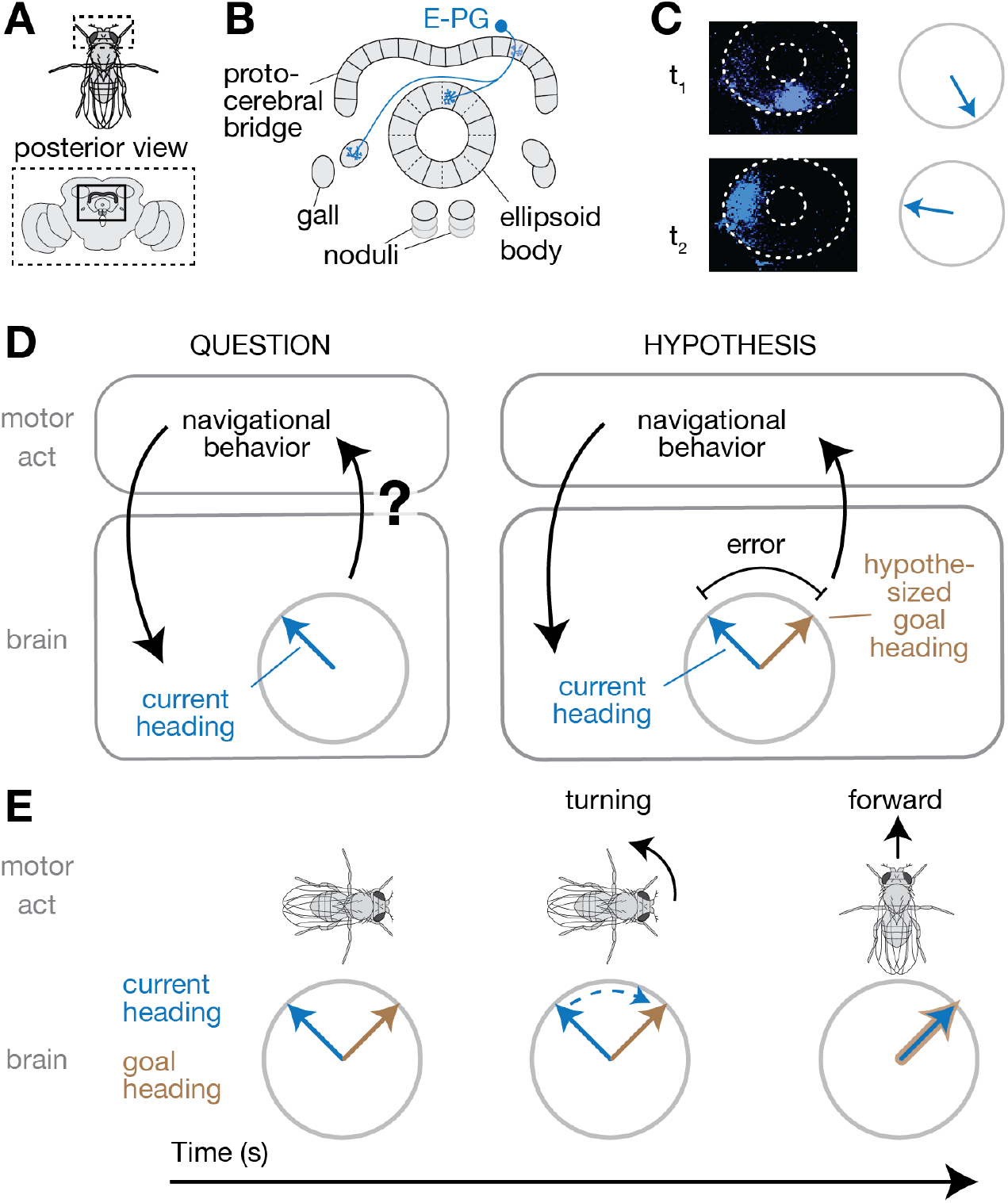
Working model for how flies compare a neural heading estimate with an internal goal angle to guide turning and walking velocities. (**A**) Posterior view of fly and brain. (**B**) Four structures in the central complex are shown, including the *protocerebral bridge* and *ellipsoid body*, with the anatomy of a single E-PG neuron indicated in blue. In each fly, an array of E-PG neurons fully tile the ellipsoid body and almost fully tile the bridge (31). (**C**) The array of E-PG neurons carries a single calcium activity peak (blue) that rotates around the donut-shaped ellipsoid body (dotted outlines) tracking the fly’s (virtual) heading (8), consistent with an internal heading estimate. Two sample frames from an E-PG>GCaMP6f imaging experiment in the ellipsoid body are displayed. E-PGs also carry 2-3 peaks that move left and right in the bridge (9, 10) (not shown). (**D**) Left: How E-PG activity impacts navigational behavior remains unknown. Right: Hypothesis that flies compare an internal heading estimate carried by E-PGs with an internal goal angle to guide their navigational behavior. (**E**) Illustration of how the model in (D) would operate to guide a fly’s behavior over time. The fly turns to align its internal heading estimate with an internal goal angle. When the two are aligned, the fly walks forward faster.

To test this model, we first verified that E-PG neurons indeed track the fly’s heading, rather than its goal heading, in our task. We reasoned that during experimental rotations of the bar, a goal signal would stay unchanged in the brain since the fly reacts to this perturbation by turning so as to bring the rotated bar back to its original position (Figure 2), arguing that the goal position was not altered by the bar jump. A heading signal, on the other hand, should update with the bar’s new location on the arena since the repositioned visual cue provides evidence to the fly that its heading has changed. We imaged E-PG activity in the protocerebral bridge under two-photon excitation (9, 18) with GCaMP6f (32) during experimentally induced, discontinuous rotations of the visual scene while the fly walked in closed-loop (Figure S4A-B). We found that the position of the periodic E-PG calcium activity peaks in the bridge, i.e. the *E-PG phase*, consistently updated with the repositioned bar, as reported in previous studies (8), and inconsistent with the E-PG phase representing the fly’s goal. Another possibility we ruled out is that the E-PG phase tracks the *difference* between the fly’s current- and goal-headings. We imaged E-PG activity in flies that changed their goal heading at least once during the course of the imaging session and found that the E-PG phase consistently followed the position of the bar on the arena, independently of where on the arena the goal was positioned, as expected from a straightforward heading signal that is ignorant of the goal position (Figure S4C-D).

If the E-PG phase represents an internal heading signal that flies compare with a goal signal to guide oriented navigation, then inhibiting E-PG neuron output should impair the ability of flies to perform this task (Figure 4A). To test this prediction, we expressed in E-PG neurons *shibire^ts^*, a dominant mutant of dynamin that impairs synaptic transmission at high temperatures (> 29°C) (33). We verified that *shibire^ts^* impaired E-PG physiology by imaging calcium activity in *shibire^ts^*-expressing E-PGs (Figure S5A). Because E-PGs show persistent activity in flies standing still in complete darkness (8, 9), it is likely that they participate in recurrent circuits that help to maintain their activity; in this case, inhibiting E-PG synaptic output should impair their own activity. Indeed, although E-PG neurons remained active when E-PG synaptic release was inhibited at 34°C, the E-PG phase was highly unstable, and poorly tracked the fly’s heading, both with a closed-loop bar (Figure S5B-D) and in the dark (Figure S5E). (We note that the E-PG phase *velocity* tracked the bar *velocity* reasonably well even in E-PG>*shibire^ts^* flies, but not well enough to prevent the heading signal from drifting wildly, see Figure S5D.) These observations show that the heading signal in E-PGs is at least partially degraded when E-PG synaptic output is impaired, and thus a behavioral feedback loop with E-PGs, if present, should also be impaired.

We measured the effect of impairing E-PG synaptic output on the flies’ walking behavior (Fig. 4). The first obvious effect was that E-PG-impaired flies dispersed less far than controls (Figure 4B). Shorter 2D dispersals would be expected if E-PG-impaired flies are worse at maintaining a consistent heading in continuous closed-loop with a bar, which indeed is what we observed (Figure 4C, Figure S6). Intriguingly, further analysis of our continuous closed-loop (Figure 4C) and bar rotation (Figure S7) experiments revealed that E-PG-impaired flies retain a residual ability to maintain the bar in front, albeit with more angular variance than the arbitrary-angle fixation seen in control flies (i.e. data points closer to the center in Figure 4C). Thus, E-PG-impaired flies show a residual form of front-bar fixation, but cannot effectively maintain their heading at an arbitrary angle relative to a visual landmark for extended periods.

**Figure 4.**
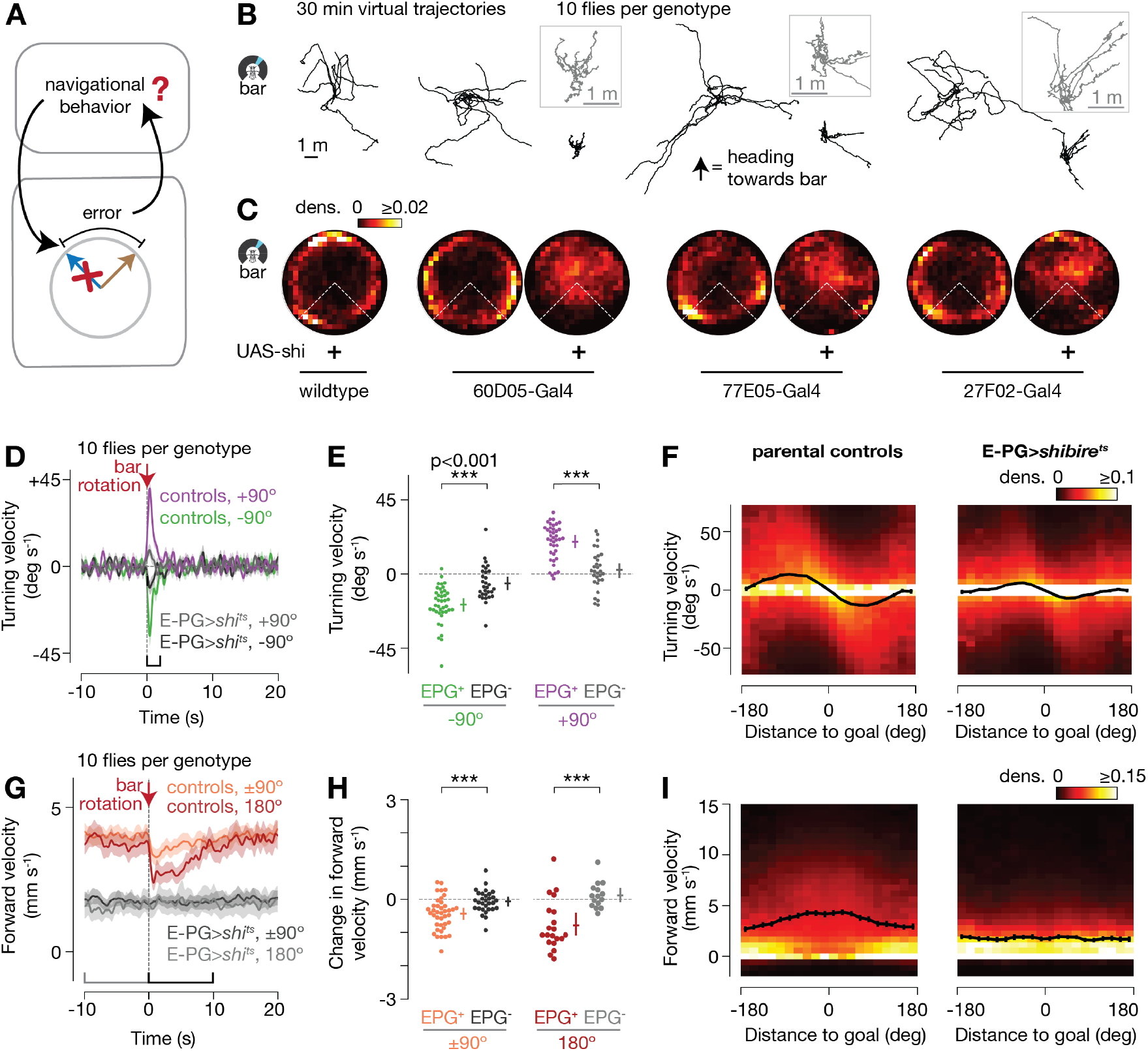
E-PG-impaired flies do not stably maintain their heading at an arbitrary angle. (**A**) How is the fly’s navigational behavior affected when E-PG synaptic output is impaired? (**B**) 2D virtual trajectories of flies walking with a bar in closed-loop. (**C**) Polar distributions of mean heading vectors from (B), taken over sliding 60 s windows (same analysis as Figure 1D). Time points where the flies were standing still (i.e. forward velocity < 0.5 mm/s)––which yield stable headings, trivially––were ignored (see *Methods*). (**D**) Turning velocity over time during 90° bar jumps. Mean and s.e.m. across flies are shown. (**E**) Mean turning velocity for each fly, computed 0 to 2 s after bar jumps. Mean and 95% confidence intervals are shown. p-values were computed using the Wilcoxon rank-sum test. (**F**) Turning velocity as a function of angular distance to goal. Data from −20 s to 40 s around bar jumps were used to create this plot. The goal was operationally defined as the mean heading averaged across the 10 s immediately prior to the bar jump. Each column of the heat map is normalized independently because we had many more data points near x=0. Mean and s.e.m. across flies are shown (black curve). (**G-I**) Same as (D-F), but for forward velocity during 90° and 180° bar jumps. In (H), we show the change in the mean forward velocity between the 10 s window immediately before and the 10 s window immediately after bar jumps for each fly. For (D-I), we only anlayzed trials where the flies maintained a relatively stable heading (circular s.d. < 45°) for 10 s before the bar jump, as an indication that flies were performing arbitrary-angle fixation. Approximately 14% and 18% of trials were excluded for control and E-PG>*shibire^ts^* flies, respectively. No forward walking requirements were applied to (D-I). For (D-I), we pooled control and E-PG>*shibire^ts^* genotypes.

E-PG-impaired flies walked more slowly than controls (Figure 4G), which also contributed to shorter dispersal distances. Indeed, the model in Figure 3 predicts slow walking in E-PG>*shibire^ts^* flies because the E-PG phase and goal are expected to be poorly matched and thus rarely drive fast forward walking in these flies (see also *Discussion*). Beyond the observation of generally slow walking, E-PG-impaired flies also showed no obvious decrease in forward walking speed following bar rotations (Figure 4G-H, Figure S8B), and a very weak (or no) relationship between forward walking speed and angular distance to goal (Figure 4I). Control flies walking at similar speeds as E-PG-impaired flies still slowed down following 180° rotations (Figure S9C-D), arguing that the lack-of-slowing-down seen in E-PG>*shibire^ts^* flies was not simply due to them walking slowly before the bar jump. E-PG-impaired flies also turned weakly in response to bar jumps (Figure 4D-E, Figure S8A), and as a function of angular distance to goal (Figure 4F). We note, however, that control flies walking at similar forward speeds as E-PG-impaired flies exhibited a similar dampening of their turning responses (Figure S9A-B), arguing that the weak turning responses of E-PG>*shibire^ts^* flies is, at least in part, due to their slow forward walking speeds. Near-wildtype turning velocities in EPG>*shibire^ts^* flies (for their walking speed) may be expected given that these flies were still able to actively, if unstably, orient toward bars in front (Figure S7), a behavior that may employ neural pathways that bypass E-PGs. These residual behavioral capacities notwithstanding, E-PG impaired flies could not stably maintain a bar on the side or in the rear, demonstrating a fundamental impairment in their ability to perform arbitrary-angle fixation. These behavioral effects were consistent across all three E-PG-expressing Gal4 lines (Figure 4B-C, Figure S5-8). Anatomical evidence argues that the only cells consistently inhibited across the three Gal4 lines are E-PGs (Figure S10 and Supplemental Table 1).

We sought to further test the model in Figure 3 via neural stimulation experiments (Figure 5A). We performed these experiments in flies walking in darkness, to assess the effects of stimulating the fly’s internal heading system independently of any impact visual inputs might have on the system. Earlier, we showed that flies do not maintain a consistent virtual heading on the ball in the dark (Figure 1D, bottom row); however, intriguingly, we observed that flies walking in darkness maintained the E-PG activity peaks in nearly as stable a position in the brain as when they walked with a closed-loop bar (Figure S11), suggesting that, under these conditions, flies walking in darkness are attempting to maintain a straight trajectory with respect to their E-PG heading signal, even if their actual trajectories on the ball drift. We experimentally rotated the E-PG phase in flies walking in darkness by activating P-ENs (<u>p</u>rotocerebral bridge-<u>e</u>llipsoid body-<u>n</u>oduli) (31), cells that are known to excite E-PGs (9, 10). (See *Methods* for why we did not stimulate E-PGs directly.) We expressed in P-ENs the ATP-gated cation channel P2X_2_, and locally released ATP on 1-2 glomeruli in the bridge (Figure 5A-C). This stimulation repositions the E-PG phase in the bridge by both activating E-PGs just medial to the stimulated location and by suppressing E-PGs (via as-of-yet uncharacterized circuitry) where they were originally active (9, 34). We measured the effect of this stimulation both on E-PG activity with GCaMP6f, and on the fly’s behavior, as it walked in the dark.

**Figure 5.**
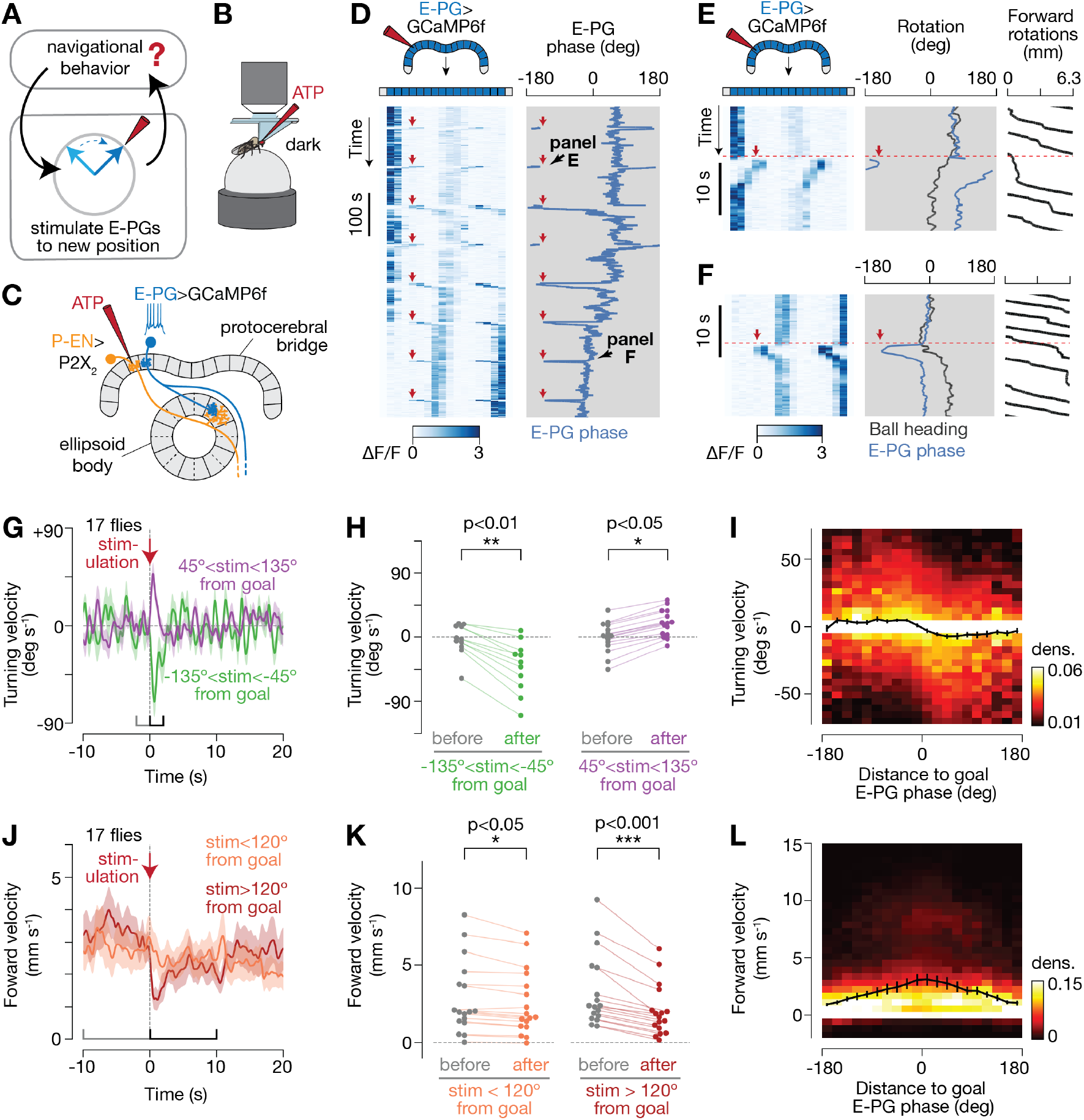
Flies slow down and turn to bring their E-PG phase back to its previous position in the protocerebral bridge after the phase is rotated via neural stimulation. (**A**) How does the fly react to a chemogenetically-stimulated change in its E-PG phase? (**B-C**) We stimulated a different central-complex cell-class, *P-ENs*(9, 10, 31), by expressing in them an ATP-gated ion channel (P2X_2_) and puffing ATP with a pipette on specific glomeruli in the protocerebral bridge, while imaging E-PGs as flies walked in the dark. Stimulating P-ENs reliably repositions the E-PG phase medial to the stimulated P-EN glomerulus (9) (Movie S1 and S2). (**D**) Sample trace where eight P-EN stimulation events reposition the E-PG phase to a consistent position in the bridge. Here, the fly gradually changed its goal E-PG phase over 12 min, as indicated by a slow drift in the mean phase signal over time. Red arrows mark stimulation times and position in the bridge, the latter of which remained constant. (**E-F**) E-PG phase, and the fly’s heading and forward walking, during individual stimulation events from (D). (**G**) Turning velocity over time during events when our stimulation was positioned 45° to 135° away from the goal E-PG phase. The goal E-PG phase was operationally defined as the mean E-PG phase in the 10 s window before stimulations. (**H**) Mean turning velocity for each fly before (−2 to 0 s) and after (0 to 2 s) stimulation. p-values were computed using the Wilcoxon signed-rank test. (**I**) Turning velocity as a function of angular distance to the goal E-PG phase. Data from −20 s to 40 s around bar jumps were used to create this plot. Mean and s.e.m. across flies are shown (black curve). Note the difference in scale of the heat map compared to Figures 2 and 4. (**J-L**) Same as (G-I), for forward velocity. In (J-K), we selected events where our stimulation was either closer (<120°) or further (>120°) away from the goal E-PG phase. In (K), forward velocities were computed over 10 s windows immediately before and after stimulations. In (G-L), we only analyzed trials where the E-PG phase was maintained at a relatively stable angle (circular s.d. < 45°) for 10 s prior to ATP stimulation, as an indication that flies were performing arbitrary-angle fixation with respect to their E-PG phase. Approximately 23% of trials were excluded. No forward walking requirements were applied to this figure.

After stimulating the E-PG phase to a new angle, we observed that flies turned on the ball in a manner that was consistent with the model in Figure 3 (Figure 5D-I). That is, they turned so as to bring the E-PG phase signal back to its location in the bridge prior to stimulation (Movie S1 and S2). The side of the bridge in which we stimulated P-ENs (left vs. right) could not explain the direction in which the flies turned (Figure S12A), and we observed no consistent behavioral turns in flies lacking a Gal4 driver (Figure S12B), arguing that these behavioral responses were caused by the rotation of the E-PG/P-EN phases (which are yoked to each other (9, 10)), specifically. In these stimulation experiments, we observed the standard relationship between the flies’ behavioral turning velocity and the angular distance to goal (of their E-PG phase) (Figure 5I), like in visual bar jump experiments (Figure 2), though the relationship seen here, during stimulation in the dark, was somewhat weaker. We also observed a decrease in forward velocity upon stimulation, which depended on the distance between stimulated- and goal-phases (Figure 5J-K). We observed the standard bell-shaped relationship between forward velocity and angular distance to goal in these experiments as well (Figure 5L). Overall, these stimulation results argue that flies turn to compensate for an experimentally-induced E-PG-phase shift in the dark, consistent with a model in which the E-PG phase interacts with a wholly internal goal angle to guide the fly’s navigational behavior, and mirroring the behavioral effects observed with rotations of visual cues on the LED arena.

## Discussion

We show that flies walking in a simple virtual environment maintain their heading at an arbitrary, goal angle (Figure 1), and actively turn back toward this angle when provided with visual evidence that they have been rotated (Figure 2). Moreover, flies tend to slow down after such rotations, and speed back up once they have corrected their heading.

Inhibiting E-PG synaptic output impaired the flies’ ability to maintain an arbitrary heading (Figure 4); however, the flies expressed a residual bias towards keeping the bar in front. This residual frontward bias may rely on visual motion- (35) or visual object- (36-39) sensitive neurons whose signals can perhaps bypass E-PGs to impact the leg motor system.

E-PG-inhibited flies walked more slowly than controls (Figure 4G). This slowing down is consistent with our model (Figure 3) because the alignment between the E-PG heading- and goal-signals is likely poor in E-PG-inhibited flies (Figure S5) and it is this alignment that we hypothesize drives fast forward walking. Alternatively, it is possible that E-PG-inhibited flies walk slowly due to some sort of general lethargy independent of any alignment between the E-PG phase and a putative goal signal. While we cannot definitively differentiate these interpretations, we note that P2X_2_-mediated stimulation of E-PGs reliably *increased* the overall level of E-PG activity in the protocerebral bridge (Figure S13), while also causing the fly to *slow down* specifically when the E-PG phase was stimulated off the goal (Figure 5J-K). This observation argues against the interpretation that E-PG-inhibited flies walk slower due to a general loss of walking drive, though more work will be needed to fully distinguish among these hypotheses.

Our data support the following working model for oriented navigation in *Drosophila*. The E-PG phase tracks the fly’s heading and is compared with an internal goal angle (Figure 3D). If the E-PG phase is positioned counterclockwise relative to the goal (when viewing the ellipsoid body from the rear), the fly tends to turn left, which in turn causes its E-PG phase to rotate clockwise, back towards the goal heading angle (Figure 3E). Conversely, if the E-PG phase is positioned clockwise relative to the goal, the fly tends to turn right. This closed-loop feedback model between a heading signal and an angular goal signal would allow the fly to maintain a steady heading, as estimated by its E-PG neurons. When the internal goal angle changes (via unknown mechanisms), the fly reorients accordingly. The fly also walks faster when its current- and goal-headings are matched, enabling the fly to travel for hundreds of body lengths along the goal direction.

The model proposed here posits the existence of angular goal signals in the fly brain; future work aimed at identifying these signals will help to further probe this model, and determine how it operates at the level of interacting neurons.

Our results focused on walking flies navigating in the context of a single, distant, vertical visual landmark. A parallel study discovered that tethered, flying *Drosophila* orient at arbitrary angles relative to a small, sun-like, dot stimulus, and that this orienting behavior is also dependent on E-PG activity (40). Future work with richer 3D visual environments will be needed to determine how the E-PG system operates with respect to local objects and the fly’s 2D or 3D location in an environment. For example, behavioral experiments have shown that fruit flies can navigate to specific locations in two-dimensional arenas (41, 42), suggesting that flies use positional, as well as angular goals, to guide behavior. Even in these tasks, however, the fly must ultimately choose a specific heading and speed at which to walk in order to reach its 2D goal. Thus, the results presented here may reflect a core interface between internal goals and motor actions, upon which other, more complex, navigational capacities in flies are built.

## Acknowledgements

We thank the Ruta and Rubin laboratories for fly stocks, Lisa Fenk for comments on the manuscript, and members of the Maimon laboratory for helpful discussions. Stocks obtained from the Bloomington Drosophila Stock Center (NIH P40OD018537) were used in this study. This work was supported by the McKnight Foundation (G.M.), the National Institutes of Health (DP2DA035148 and R01NS104934) (G.M.) and the Leon Levy Foundation (V.V.).

## Author Contributions

J.G., V.V., and G.M. conceived of the project. J.G. performed all two-photon imaging and stimulation experiments, P. M.-P. performed all purely behavioral experiments in wildtype and *shibire*-impaired flies. J.G., V.V., P.M.-P. and G.M. analyzed and interpreted all experiments. V.V. developed the bar jump perturbation approach for studying goal-directed walking. A.A. performed and analyzed all multi-color-flip-out anatomical experiments. J.G. and G.M. wrote the paper.

## Supplementary Materials

**Figure S1.**
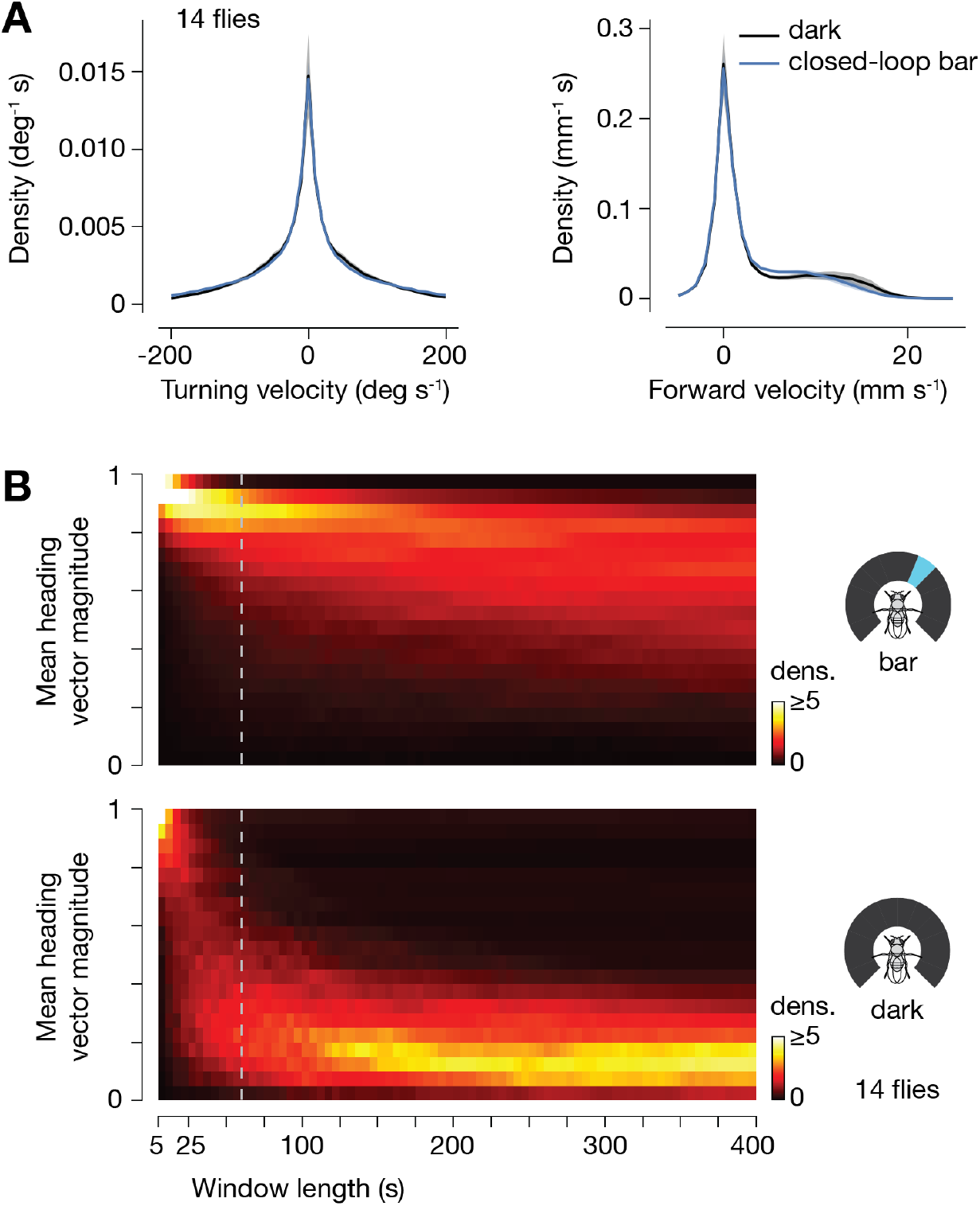
Flies maintain their heading along a relative stable angle when walking in closed-loop with a bar, but not in the dark, even though they turn and walk forward with similar first-order statistics in both conditions. (**A**) Distribution of turning and forward walking velocities in closed-loop bar and dark conditions. The mean and s.e.m. across flies are shown. (**B**). Distribution of mean heading vector magnitudes computed over different window lengths. Dashed line indicates the 60 s window used to compute the polar plots in Figure 1D. Like in Figure 1D, time points in which flies were standing still (i.e. forward velocity < 0.5 mm/s) were ignored for the calculation of mean heading vectors because heading values during such time points are stable for the trivial reason that the fly is not moving.

**Figure S2.**
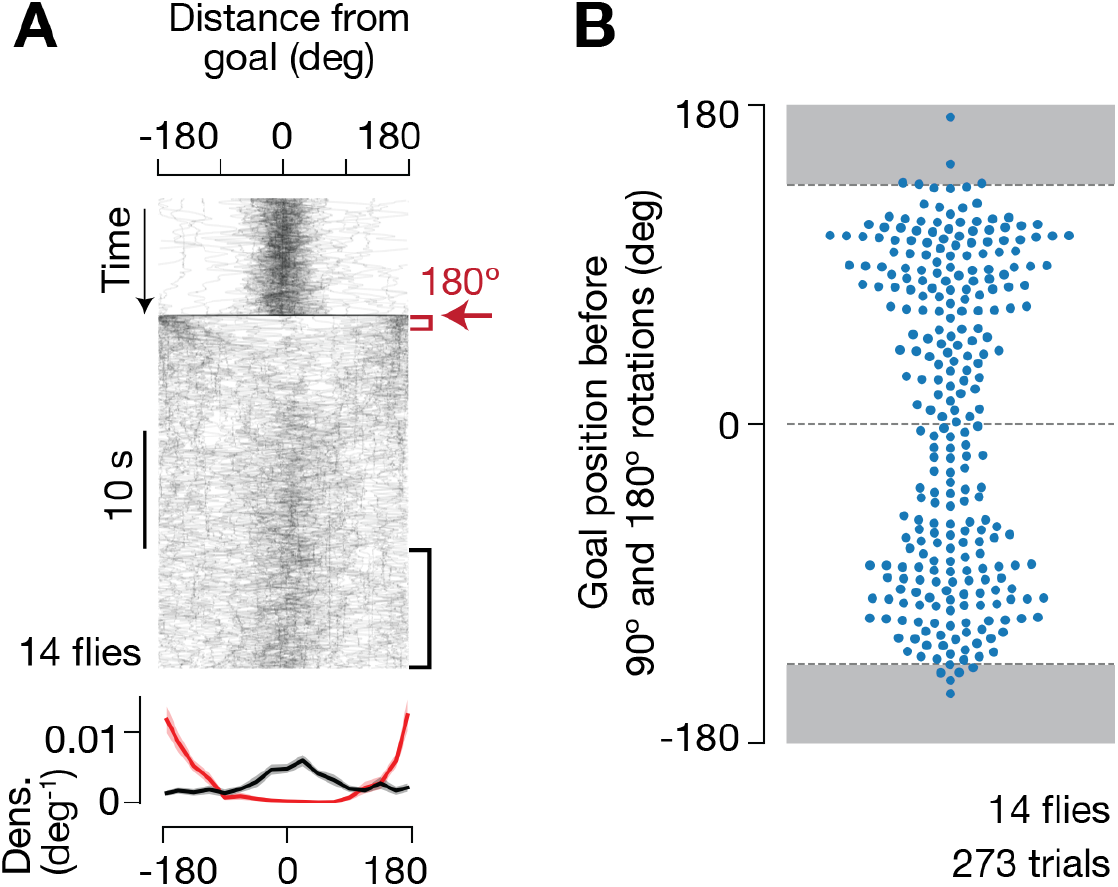
Flies turn to bring a bar to the previous (goal) heading after 180° bar jumps and goal heading angles before 90° and 180° bar jumps are broadly distributed from 0-360°, with a bias to the rear-right and rear-left of the LED display. (**A**) Top: Distance from goal over time for 180° bar jumps. The goal was operationally defined as the mean heading in the 10 s window immediately before bar jumps. Red arrow indicates when the 180° bar jump occurred. Bottom: Distance from goal distributions 0-1 s (red) and 20-30 s (black) after bar jumps. Mean and s.e.m. across flies are shown. (**B**) Distribution of goal headings (as defined above) before 90° and 180° bar jumps. In (A-B), we included all trials where the fly maintained its heading at a relatively stable location (circular s.d. < 45°) for 10 s before the bar jump. Approximately 10% of trials were excluded. No forward walking requirements were applied to this figure.

**Figure S3.**
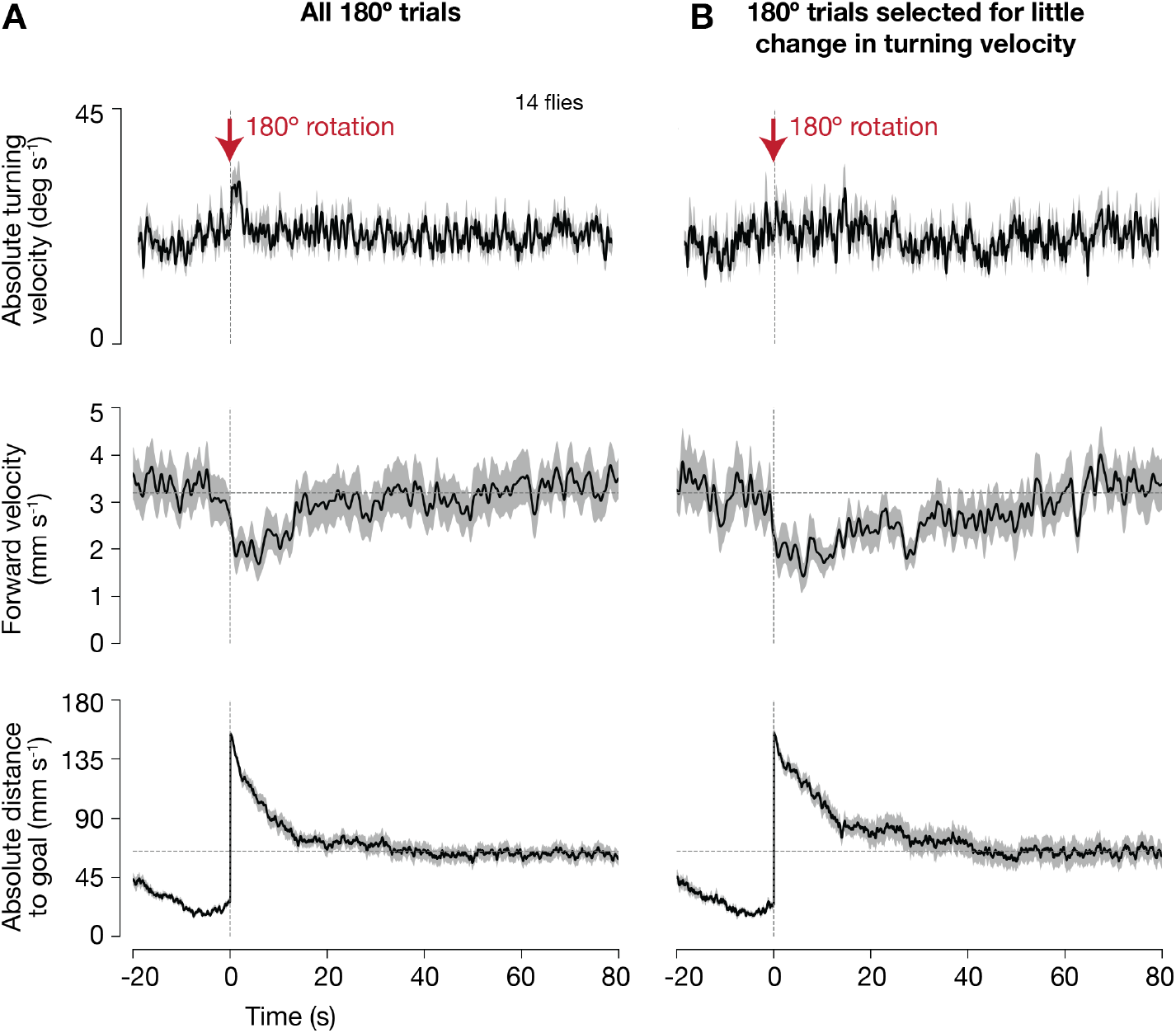
Slowing down of forward speed after bar rotations is a separable behavioral response from strong turning. (**A**) Turning and forward walking behavior of flies during 180° bar rotations. Absolute distance to goal is shown on the bottom. Time zero indicates when the bar jumped. All trials are included. Mean and s.e.m. across flies are shown. Absolute turning velocity is shown in the top plot because flies are equally likely to turn left or right after 180° jumps. These data show that flies turn and slow down after 180° bar jumps. Note that the distance-to-goal after the jump does not return to its baseline level before the jump (bottom plots) for several potential reasons: (1) the bar sometimes ends up invisible when jumped to the rear of the arena, where there are no LEDs (and the fly may take a very long time to correct this specific perturbation), (2) flies might be in a different behavioral state (grooming, sleeping) when the bar is jumped and thus may take longer than 80 s to correct the perturbation, (3) flies may sometimes change their goal heading after a bar jump (perhaps as a result of the bar jump), and (4) it is possible that prior to some bar jumps, the bar was located off the actual, internal, goal angle, which would make our operational definition of the goal inaccurate, leading flies to not return the bar to our operationally defined goal angle. (**B**) Same as (A), but selected for trials in which the flies’ turning speed changed very little in either direction (< 20°/s change in either direction comparing a 5 s window before and after the bar rotation). Flies still slowed down on these low-turn-velocity trials (middle). Since the flies turned very weakly, they returned the bar over a longer timescale, as evidenced by the slower decay of their absolute distance to goal over time (bottom), which also matched the longer timescale of their slowing-down response.

**Figure S4.**
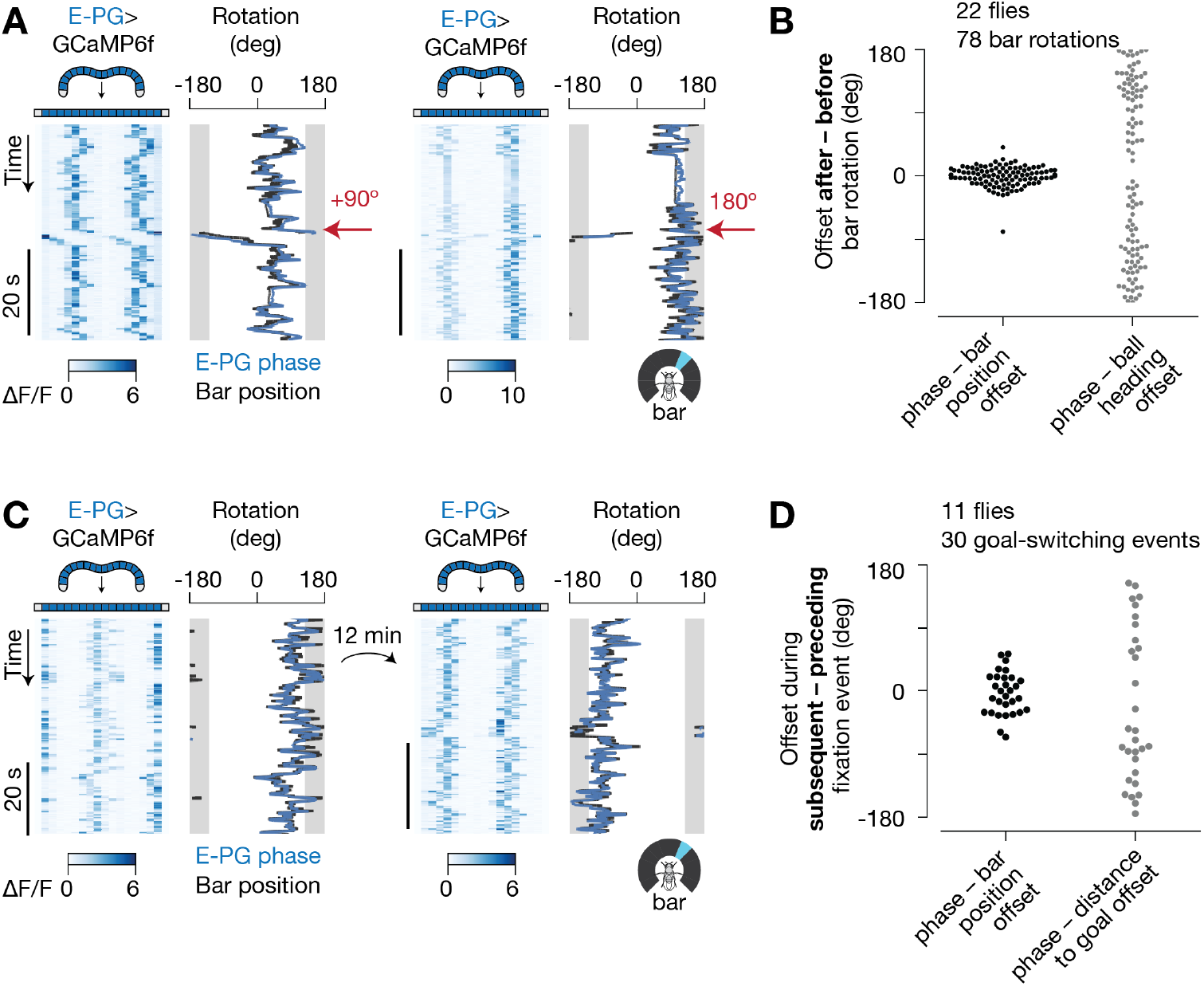
The position of the E-PG peaks in the protocerebral bridge (E-PG phase) does not track the fly’s angular goal, or the fly’s angular distance to goal, as defined in this task. (**A**) E-PG activity during 90° (left) and 180° (right) bar jumps, in two different flies (GCaMP data are analyzed as in ref. 9; see *Methods*). The 90° gap in the back of the LED display, where the bar is not visible, is highlighted in grey. To aid a visual comparison between E-PG phase and bar position, we applied a fixed offset to the E-PG phase trace (blue line), chosen to minimize offsets with the bar position (black line) over time. This is a standard transformation (8,9) since there is a variable offset fly-to-fly between E-PG bolus positions in the brain and bar positions on the LED display. We applied a different offset to the traces shown from these two flies. The distribution of E-PG phase-to-bar position offsets is roughly uniform across flies (8). (**B**) Change in offset between E-PG phase and bar position, as well as between E-PG phase and ball heading, after bar jumps. We included all trials where the fly maintained a stable heading (circular s.d. < 45°) for 10 s before the bar jump, indicating that flies are actively fixating, although the same results hold if all trials are included ((8), data not shown). The fact that the offset with respect to the bar stays relatively constant after a bar jump, but not the offset relative to the ball angle, indicates that the E-PG phase tracks the flies’ current heading (as indicated by the bar) rather than its goal heading. (**C**) E-PG activity before and after the fly changed its goal heading. Here, the same offset was applied to the traces on the left and the right, which represent data acquired from the same fly 12 min apart. (**D**) Change in offset between E-PG phase and bar position, as well as between E-PG phase and angular distance to goal, after flies changed goal headings (see Methods). That the offset between the E-PG phase and bar position does not change in either of these two conditions indicates that the E-PG phase tracks the fly’s orientation in reference to the bar, rather than other variables related to the goal. See text for details.

**Figure S5.**
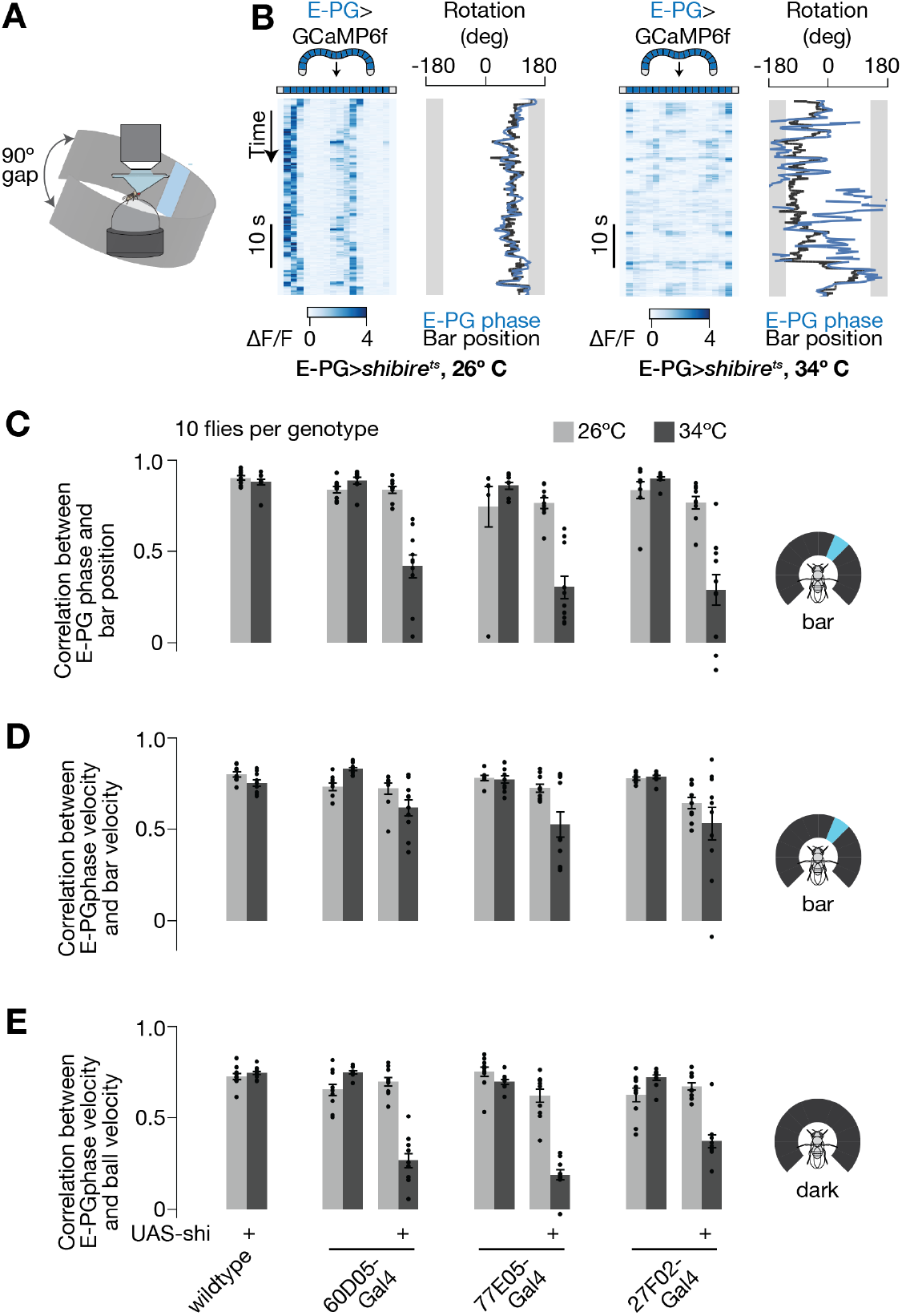
Blocking E-PG synaptic output impairs the ability of the E-PG phase both to track the flies’ heading in closed-loop with a bar and the flies’ turning velocity in the dark. (**A**) We imaged E-PG activity as the fly walked on a ball in closed-loop with a bar and in the dark (not depicted). (**B**) E-PG activity in the protocerebral bridge with E-PGs expressing *shibire^ts^* at 26°C and at 34°C. The 90° gap in the back of the arena where the bar is not visible is highlighted in grey. (**C**) Pearson correlation coefficients between E-PG phase and bar position. (**D**) Pearson correlation coefficients between E-PG phase velocity and bar velocity. These correlation coefficients are not greatly reduced by expressing *shibire^ts^* in E-PGs, suggesting that visual-motion inputs can still induce properly signed rotations of the E-PG heading signal in E-PG>*shibire^ts^* flies; however, the fidelity of such velocity updating is not sufficient to prevent large drifts over time in the angular position of the E-PG phase relative to the angular position of the bar (C). (**E**) Same as (D), but in the dark. Mean and s.e.m. across flies are shown. The cold-to-hot changes in correlation in (C) and (E) were all significantly different between E-PG>*shibire^ts^* and control groups (P < 0.01, after correcting for multiple comparisons, Wilcoxon rank-sum test). The changes in correlation in (D) were not significant.

**Figure S6.**
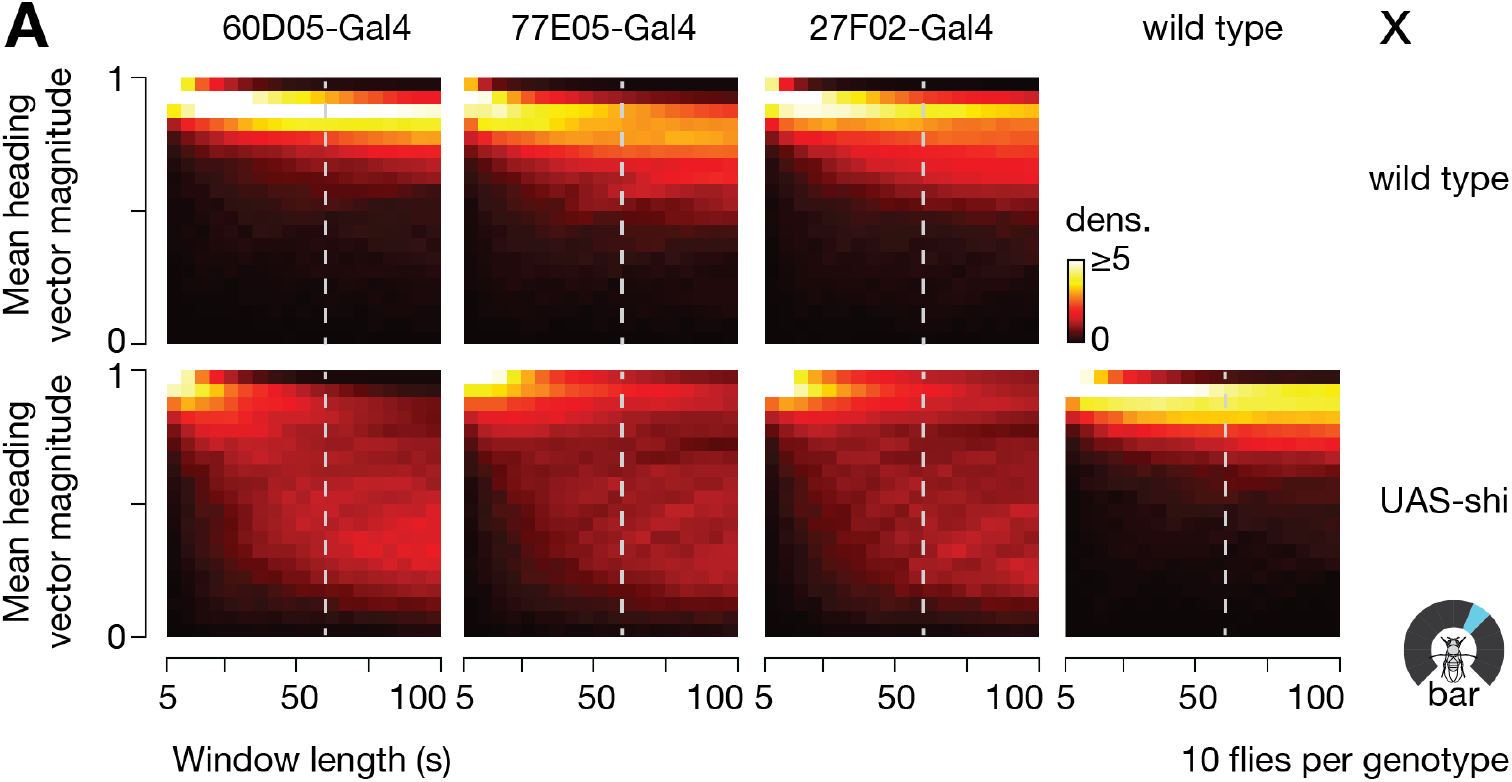
Blocking E-PG synaptic output impairs the fly’s ability to maintain its heading over multiple timescales. (**A**) Distributions of mean heading vector magnitude for different window lengths, across E-PG>*shibire^ts^* and control genotypes. The dashed line indicates the 60 s window length used in generating the polar plots in Figure 4C. Like in Figure 4C, time points in which flies were standing still (i.e. forward velocity < 0.5 mm/s) were ignored for the calculation of mean heading vectors because heading values during such time points are stable for the trivial reason that the fly is not moving.

**Figure S7.**
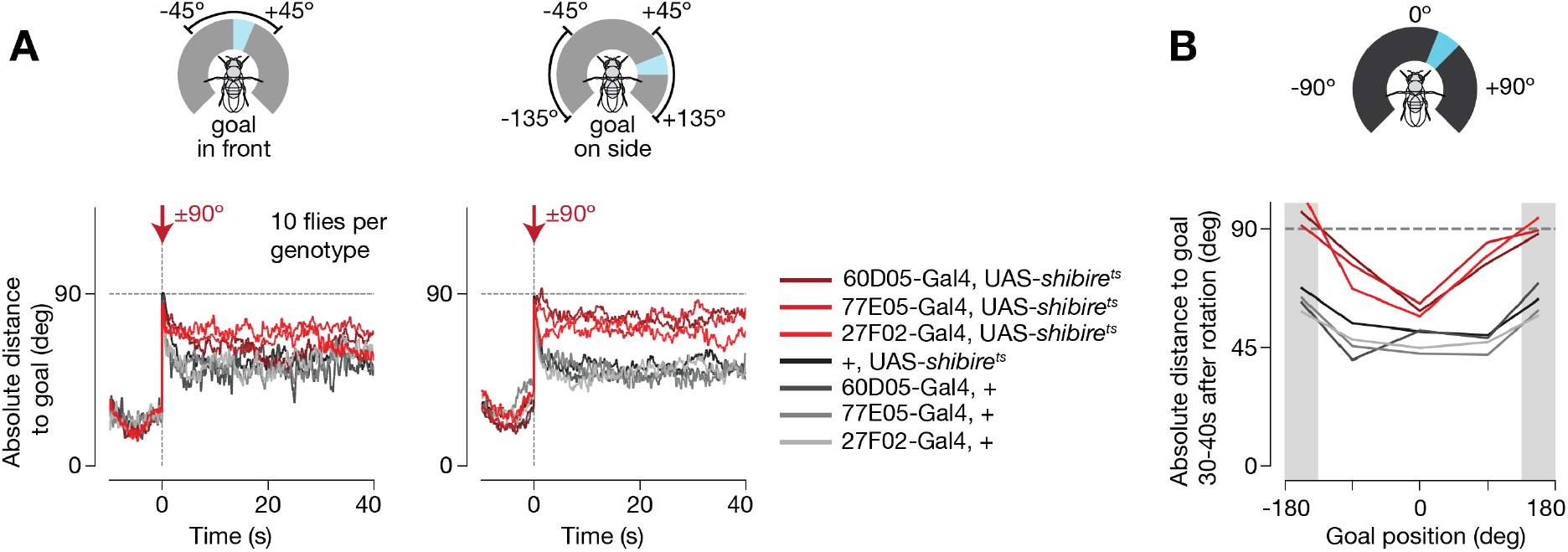
E-PG-impaired flies can correct for bar jumps in front, but less so in other visible parts of the arena. (**A**) Mean absolute distance to goal before and after 90° bar jumps, when the goal position was with the bar in front (left), or in the remaining visible parts of the arena (right), across genotypes. The fact that the gray curves come down after the 90° jump indicates that control flies perform an active correction toward the initial goal position on many trials. The fact that the red curves come down when the bar is in front (left), but not as much when it is on the sides (right), indicates that E-PG>*shibire^ts^* flies can turn in the proper direction to correct for a bar jump in front, but their behavior is less effective or ineffective when the bar is on the sides. See Figure S3 legend for a detailed explanation on why none of the curves, even the control ones, return to baseline after the bar jump. (**B**) Mean absolute distance to goal 30-40 s after 90° bar jumps, across goal positions. Control flies correct for bar jumps, irrespective of goal position, provided the goal is visible (as evidenced by a low flat line across goal positions). E-PG>*shibire^ts^* flies, however, are more impaired in their ability to correct for bar jumps the further their goal is from the front. Grey areas highlight goal positions where the bar is not visible.

**Figure S8.**
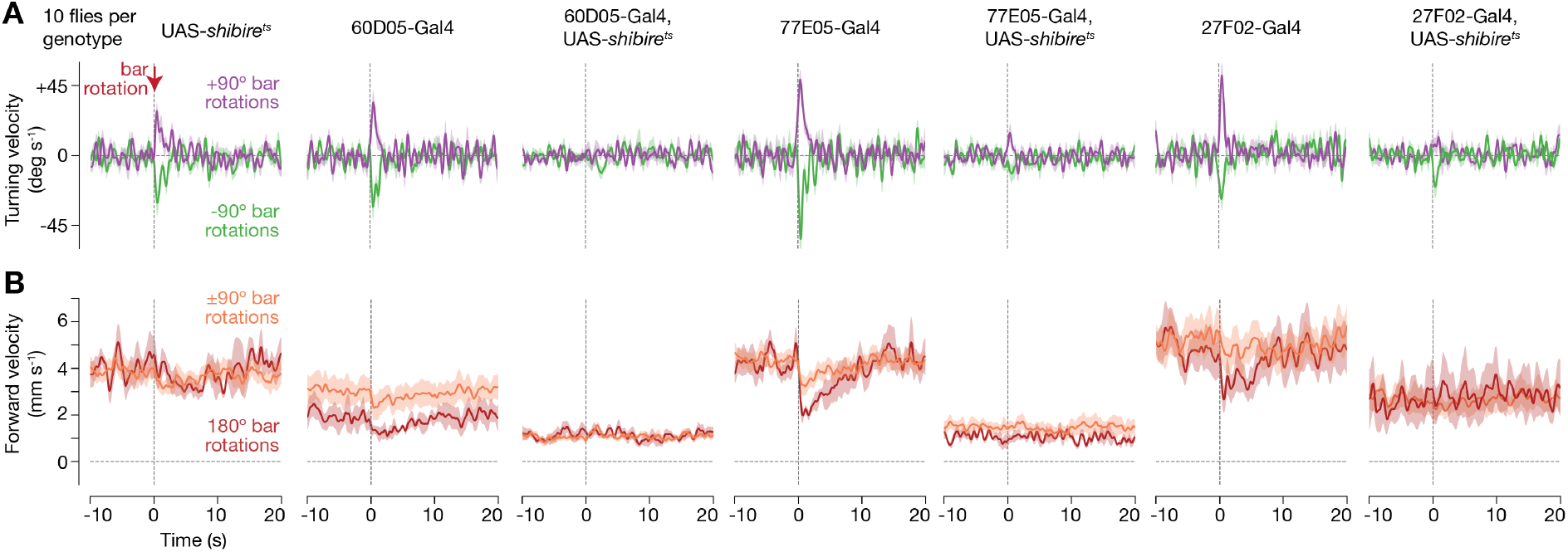
Blocking E-PG synaptic output impairs the fly’s ability to behaviorally respond to bar jumps. (**A**) Mean turning velocity over time during 90° bar jumps, across control and E-PG>*shibire^ts^* genotypes. (**B**) Same as (A), for forward velocity during 90° and 180° bar jumps. Mean and s.e.m. across flies are shown in all panels. No forward walking requirements were applied in this figure.

**Figure S9.**
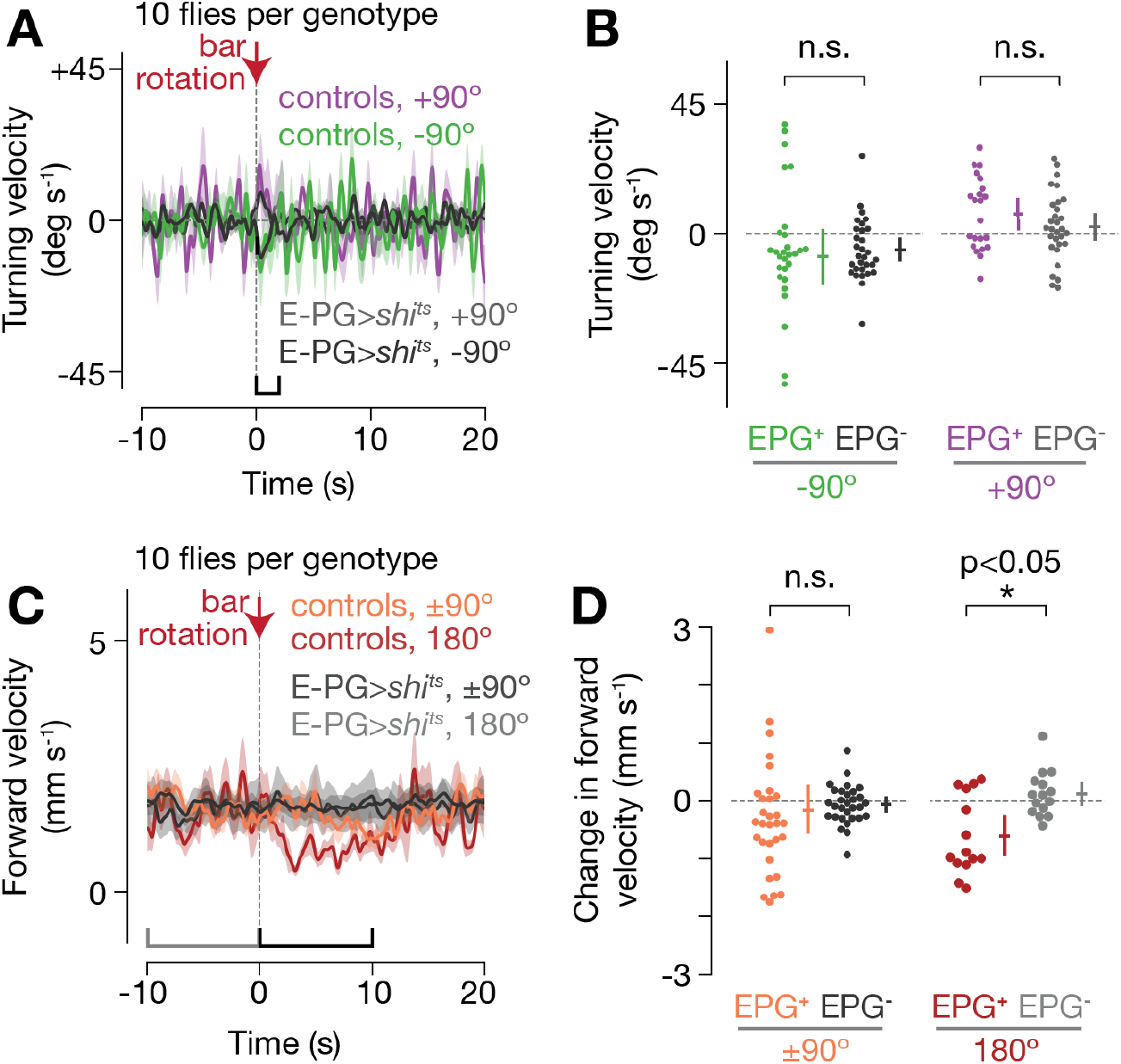
E-PG>*shibire^ts^* flies do not slow down their forward speed after bar jumps even though control flies matched for their overall forward walking speed do slow down. Data plotted as in Figure 4D-E, G-H, except selecting for trials in control flies where the flies walked––on average over the 10 s window before the bar jump––as slow as E-PG>*shibire^ts^* flies. Data from E-PG>*shibire^ts^* flies are plotted identically as in Figure 4D-E, G-H. (**A**) Turning velocity over time during 90° bar jumps. Mean and s.e.m. across flies are shown. (**B**) Mean turning velocity for each fly, computed 0 to 2 s after bar jumps. Mean and 95% confidence intervals are shown. p-values were computed using the Wilcoxon rank-sum test. (**C-D**) Same as (A-B), but for forward velocity during 90° and 180° bar jumps. In (D), we plot the change in the mean forward velocity in the 10 s window immediately before the bar jump compared to the 10 s window immediately after the bar jump, for each fly. For all panels, we included all trials where the flies maintained a relatively stable heading (circular s.d. < 45°) over the 10 s window before the bar jump, as an indication that flies were performing arbitrary-angle fixation. Approximately 78% and 18% of trials were excluded for control and E-PG>*shibire^ts^* flies, respectively. (A large number of trials were excluded in control flies because we were focusing, in these control analyses, on slow walking trials, which were rare in controls.) Four control and three E-PG>*shibire^ts^* genotypes were pooled for these analyses.

**Figure S10.**
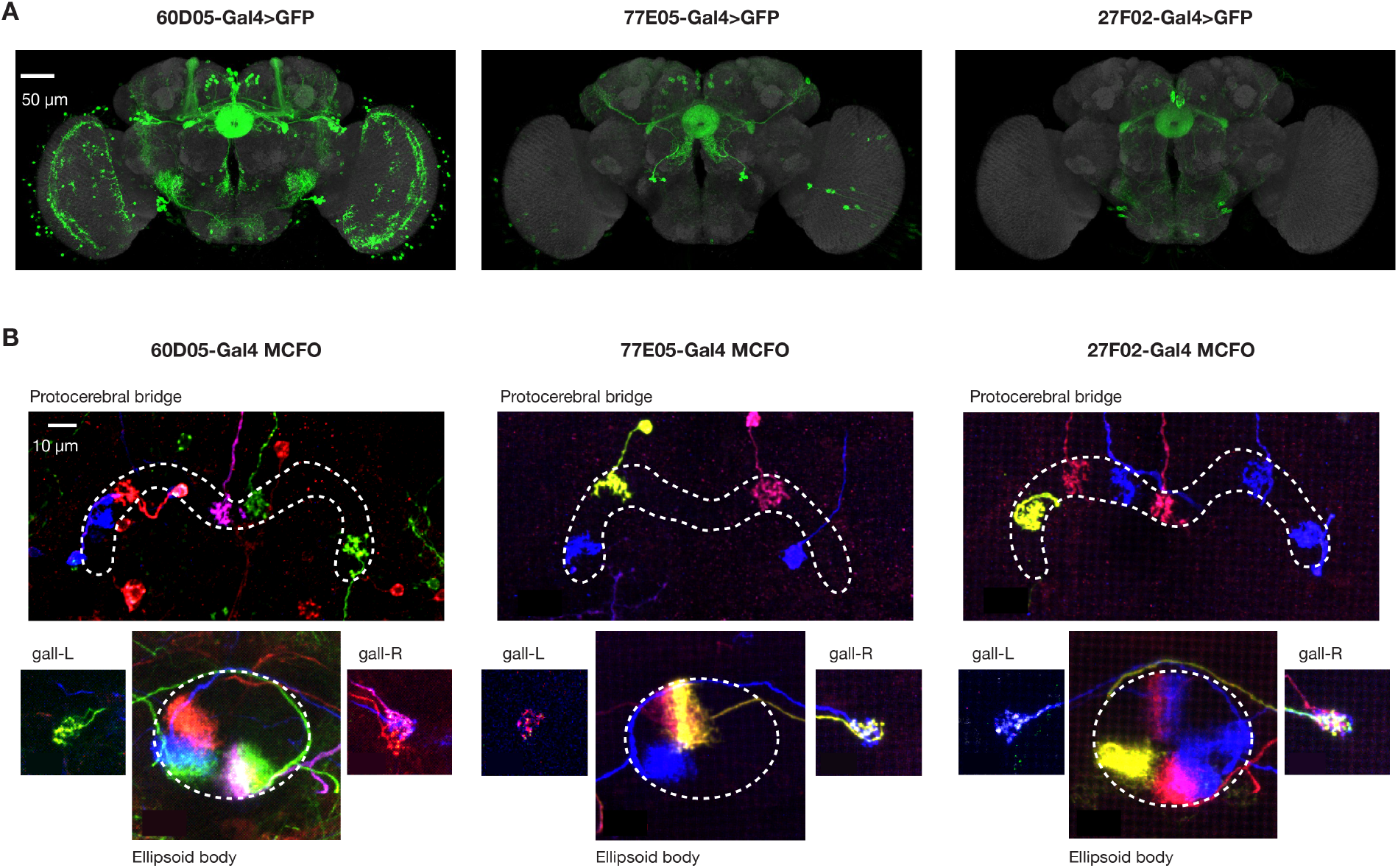
Three Gal4 lines label E-PG neurons, specifically, in the protocerebral bridge and ellipsoid body. **(A**) Maximum z-projections of three Gal4 lines driving mCD8:GFP. GFP is in green, neuropil (nc82) is in grey. The only common cell type clearly labeled by these three Gal4 lines is E-PGs in the central complex. Data were downloaded from http://flybrain.mrc-lmb.cam.ac.uk/ (43) and are reproductions of publically available Gal4 expression data (44). (**B**) Three example brains, one per Gal4 line, in which neurons were stochastically labeled in different colors by the multicolor flip-out (MCFO) method (45). E-PG neurons, which innervate the ellipsoid body, protocerebral bridge and gall, was the only cell type we observed in the central complex. 46 of 48 MCFO-visualized neurons that innervated the central complex in these three Gal4 lines were unambiguously E-PGs (and all 48 might have been E-PGs) (see Supplemental Table 1 for the entire dataset).

**Figure S11.**
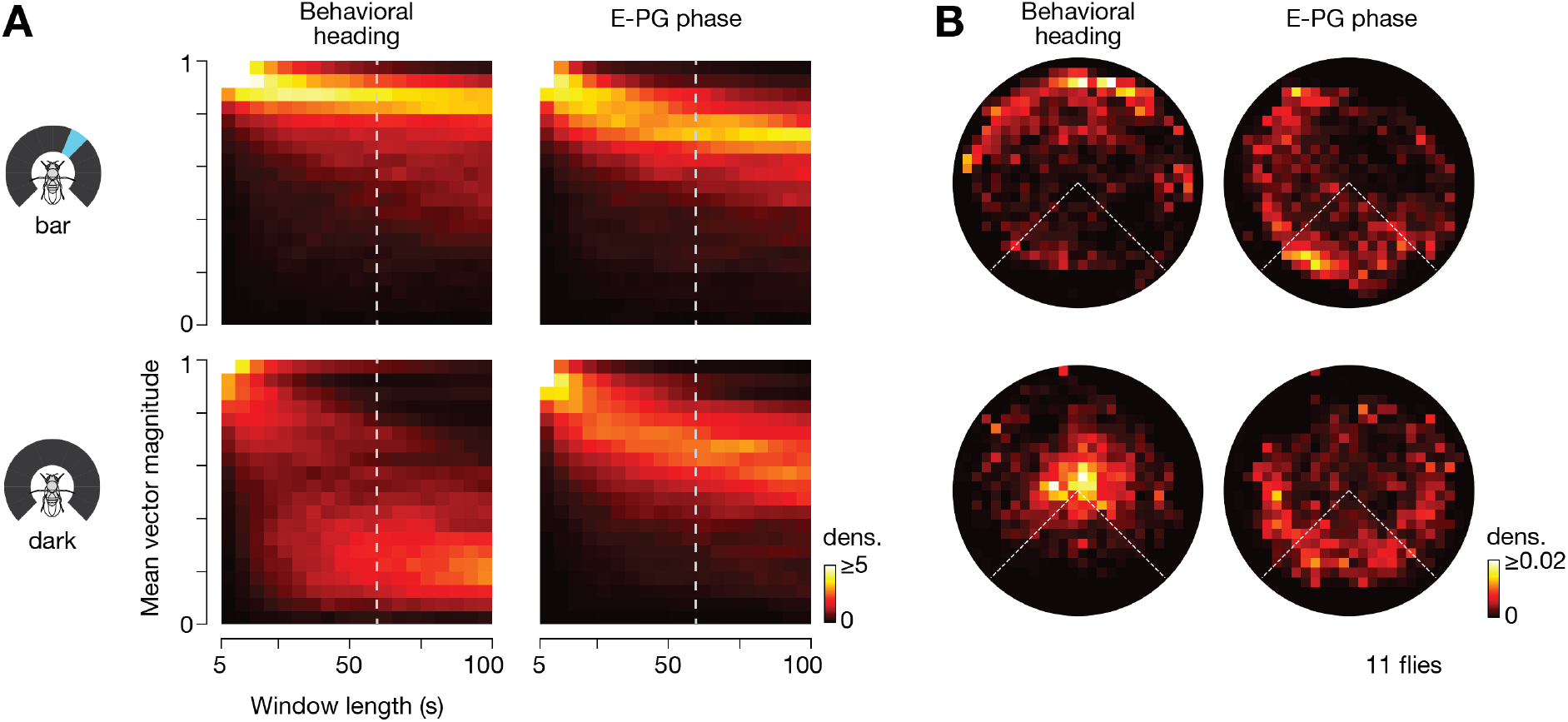
In complete darkness, even though the 2D trajectories of flies drift over time, the E-PG heading signal is maintained at a relatively stable position in the brain, suggesting that flies are still attempting to use their E-PG signal to walk straight. (**A**) We imaged E-PG activity over 10 min of closed-loop bar, and 10 min of constant darkness (the order was reversed in half the flies). Shown are the mean vector magnitude distributions computed over different window lengths for the fly’s heading on the ball and E-PG phase in the brain during closed-loop bar and dark conditions. Dashed line indicates the 60 s window length used to generate the polar plots in B. (**B**) Polar distributions of mean vectors taken over sliding 60 s windows of walking, for the fly’s heading and E-PG phase in closed-loop bar and dark conditions. The fact that the two plots on the right of (A) and (B) look similar argues that flies maintain their E-PG phase at a consistent, goal, location while walking in complete darkness. Their behavioral heading, however, drifts in darkness (bottom left plots in (A) and (B)) because the flies have no visual feedback to inform them that their walking movements are causing them to go off course.

**Figure S12.**
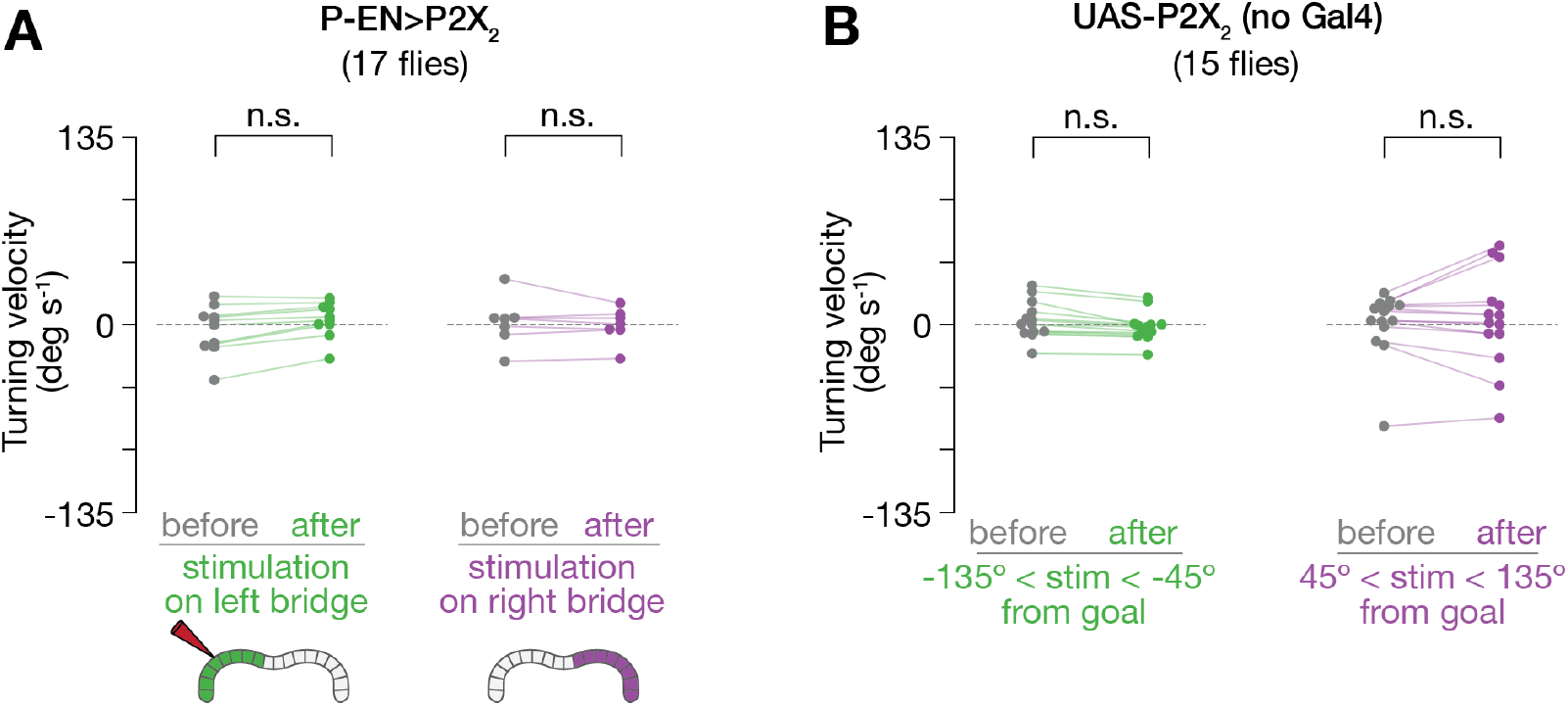
On average, flies do not turn in response to chemogenetically stimulating P-ENs in the left or right bridge, or if P2X_2_ expression is not driven by Gal4. (**A**) Mean turning velocity in each fly before (−2 to 0 s) and after (0 to 2 s) stimulating the left and right bridge, in flies expressing P2X_2_ in P-ENs. These data show that the side of the bridge in which we stimulate P-ENs does not predict which way the flies turn. Rather, the rotational direction of the E-PG phase caused by stimulation of P-ENs (which varies from trial to trial and is independent of which side of the bridge we stimulate) is the predictive variable (Figure 5), consistent with our model (Figure 3). (**B**) Turning velocity in flies lacking a Gal4 driver before (−2 to 0 s) and after (0 to 2 s) events when the ATP pipette was positioned 45° to 135° away from the goal E-PG phase, as in Figure 5H. Like in Figure 5, the goal E-PG phase was operationally defined as the mean E-PG phase in the 10 s window immediately before ATP release. There is no measureable effect on turning velocity when puffing ATP onto the protocerebral bridge in a fly that does not express P2X_2_ in P-ENs. Statistical comparisons were computed using the Wilcoxon signed-rank test.

**Figure S13.**
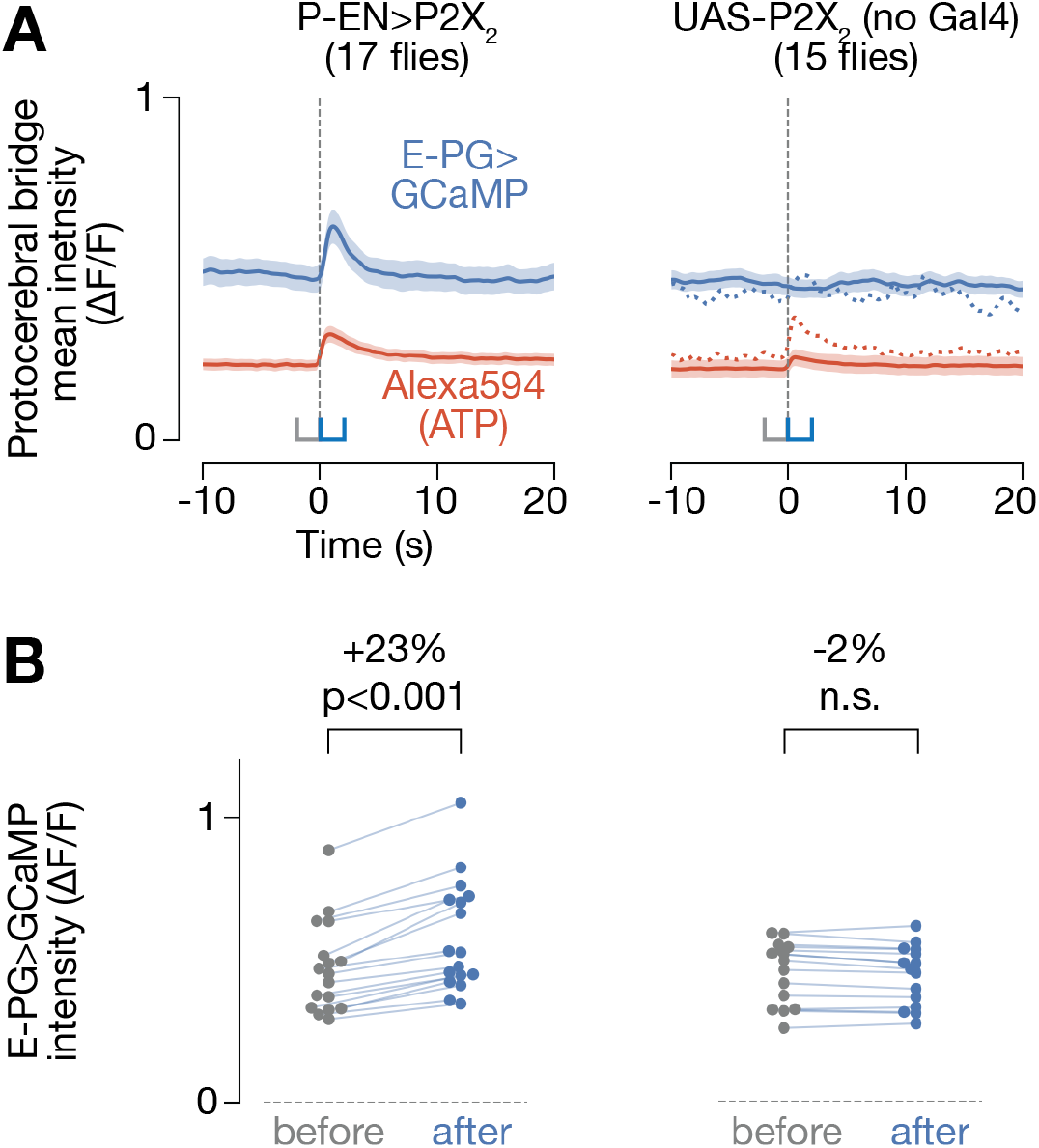
Stimulating P-ENs causes an overall increase in E-PG activity. (**A**) Mean fluorescent intensity across the entire protocerebral bridge over time during ATP stimulations. Blue trace represents the E-PG>GCaMP signal. Red trace represents the Alexa594 signal. Alexa594 was co-loaded with ATP in the pipette to visualize the release of ATP. Mean and s.e.m. across trials are shown. We observed an increase in E-PG GCaMP in experimental (P-EN>P2X_2_) (left) but not control (UAS-P2X_2_;no Gal4) (right) flies. For unclear reasons, the Alexa-594 signal was weaker during stimulation of control flies than during stimulation of experimental flies, suggesting less ATP may have been applied to the bridge. We therefore selected for trials in control experiments in which the mean increase in the Alexa594 signal was similar to the mean increase measured in non-controls and we still observed that the mean E-PG GCaMP signal did not measurably increase during ATP puffs. (**B**) Mean E-PG>GCaMP intensity for each fly before (−2 to 0 s) and after (0 to 2 s) stimulation. p-values were computed using the Wilcoxon signed-rank test. In (A-B), we only included trials where the E-PG phase was maintained at a relatively stable location (circular s.d. < 45°) in the 10 s window immediately before stimulation, to match the data used in this analysis with that used in Figure 5.

## Materials and Methods

### Fly stocks

Flies were raised with a 12 hour light, 12 hour dark cycle. All experiments, except immunohistochemistry experiments, were performed with 1-3 day old females with at least one wild-type white allele. Flies were selected randomly for all experiments. We were not blind to the flies’ genotypes. Genotypes and their origins are listed in Supplemental Tables 2, 3.

### Immunohistochemistry

Dissection of fly brains, fixation, and staining for neuropil and multicolor flip-out antigens were performed as previously described (9, 45).

### Behavioral setup

Tethered walking, behavioral imaging, ball tracking, and closed-loop visual feedback were setup as previously described (9). We used different rigs for our behavioral and imaging experiments. For the behavioral rig, we recorded data on a National Instruments BNC-2090A A/D converter. For the imaging rig, we recorded data on an Axon Instruments Digidata 1440 A/D converter. Otherwise the two rigs were as identical as possible, including matching the fly’s position relative to the LED display (arena) and the angle of the LED display (see Behavioral task below). We recorded behavioral data at 50 Hz. The temperature of the fly’s head and upper thorax were controlled by the temperature of water (or physiological saline in the case of imaging experiments) flowing over these body regions.

### Behavioral conditions

Flies were starved for 8-16 hours prior to testing, and were heated via the water or saline bath to at least 30°C (measured in the bath), except for E-PG>*shibire^ts^* control imaging experiments, where we kept the bath at 26°C (Figure S5). We originally used these conditions to promote walking; however, we also found that the flies tended to maintain their heading at a consistent, but arbitrary angle with respect to the bar under these conditions. We note that these experimental parameters likely reflect an uncomfortable situation for the fly, and that the fly is likely reacting by attempting to disperse, or search a large area in search of food or cooler temperatures. In addition, the flies’ heads were glued to our fly plates with a downward pitch (antennae angled down, toward the ball), since this posture facilitated protocerebral bridge imaging and ATP stimulation with a pipette. We kept this posture consistent across all imaging and behavioral experiments (always using physiology plates (19), not pins, for experiments). To partially account for this pitch, we tilted the arena 30° forward with respect to the horizontal (angled arena in Figure 1A), to better match the angle of the fly’s head. However, it is likely that the flies still viewed the bright bar with their dorsal visual field, which might lead them to interpret our bright visual landmark (see below) as a celestial cue, useful for orienting their dispersal.

### Visual stimuli

We used a cylindrical LED arena (20), spanning 270° in azimuth, and 81° in height. However, we note that the ball below the fly and the physiology plate above the fly’s head acted as visual occluders, allowing the flies to see only ~45-50° of the full height of the bar. For closed-loop bar experiments, we presented the fly with a 6 pixel-wide (11.25°), 81° high bright bar. The bar rotated with the ball with a gain of 1. For discontinuous rotations of the visual stimulus, the fly was presented with a closed-loop bar, except for one frame every trial where the bar jumped −90°, +90° or 180° from its current position. Immediately after the bar jump, it resumed rotating in closed-loop with the fly from its new position. For the 30 s dark stimulus, the fly was presented with a closed-loop bar for 3 minutes, then 30 s of constant darkness, after which the bar reappeared with a random offset with respect to the ball. For all experiments with a closed-loop bar, we let the fly walk in closed-loop with the bar for at least 5 minutes before beginning each experiment. Trial orders were randomly permuted for all experiments. See Supplemental Table 2 for trial structure for each experiment.

### Imaging setup

Calcium imaging, two photon data acquisition and alignment with behavioral data were performed as previously described (9). We excited GCaMP6f with a Chameleon Ultra II Ti:Sapphire tuned to 925 nm, with 20-50 mW at the back aperture. We recorded all imaging data using 3-6 z-slices at a volumetric rate of 4-6 Hz. In all figures, the left protocerebral bridge is shown on the left, and the right bridge on the right. In Movie S1 and S2, this orientation is flipped, since the fly is viewed from the front. We perfused the brain with extracellular saline composed of, in mM: 103 NaCl, 3 KCl, 5 N-Tris(hydroxymethyl) methyl-2-aminoethanesulfonic acid (TES), 10 trehalose, 10 glucose, 2 sucrose, 26 NaHCO_3_, 1 NaH2PO_4_, 1.5 CaCl_2_, 4 MgCl_2_, and bubbled with 95% O_2_ / 5% CO_2_. The saline had a pH of 7.3-7.4, and an osmolarity of 280±5 mOsm.

### P2X2-based stimulation

The most straightforward way to reposition the phase of the E-PG signal in the protocerebral bridge would be to stimulate E-PGs directly. Although we could reliably stimulate E-PGs by expressing P2X_2_ in them directly (data not shown), we were not able to induce bilateral changes in E-PG activity in the bridge by stimulating one position on one side of the bridge with stimulation strengths that elicited E-PG responses within their physiological range. However, we could regularly achieve a naturalistic, bilateral, re-positioning of the E-PG activity peaks via stimulation of a different cell class in the central complex, called P-ENs, at one location in the bridge (Figure 5). P-EN neurons from one glomerulus on one side of the bridge project to a tile in the ellipsoid body that contains E-PG neurons that project to the left bridge as well as E-PG neurons that project to the right bridge (31), which may explain why P-EN stimulation in the bridge is more effective at repositioning E-PG activity bilaterally. We note that others were able to relocate the E-PG activity peak by directly activating E-PGs (via optogenetic stimulation) in the ellipsoid body (34), which is innervated by E-PGs from both sides of the bridge. We preferred to chemogenetically activate E-PGs (via P-ENs), since we found that optogenetic light stimulation tended to cause the fly’s behavior to change (including turning), irrespective of whether we expressed a light-activated channel in any neuron. Thus, we stimulated E-PGs by chemogenetically exciting P-ENs. Specifically, we expressed in P-ENs the ATP-gated cation channel P2X2, and focally released ATP from a pipette via pressure pulses, as described previously (9). Unlike previous experiments, however, here we analyzed the fly’s behavioral responses to stimulations and provided ATP pulses with longer inter-pulse intervals (2 min.) to ensure that flies had ample time to behaviorally correct for each perturbation. We used 0.5 mM ATP in extracellular saline delivered with 20-50 ms pressure pulses. To compute the stimulated E-PG phase, we computed the mean E-PG phase over 0.5 to 1 s after stimulations for each recording. We collected data from 17 experimental and 15 control flies.

### Data analysis

Two photon images over time were first registered along the x-y plane using python 2.7, as described previously (9). Regions of interest highlighting the 18 glomeruli in the bridge were then manually parsed in Fiji (9, 46) and all subsequent analyses of these signals were performed in python 2.7 as described previously (9). Some flies, which seemed excessively starved and appeared quite unhealthy as a result, were not analyzed; otherwise, no flies were excluded. No statistical method was used to choose the sample size. When plotting turning or forward velocity over time (e.g. Figure 2, 4, 5), these signals were smoothed with a 200 ms Gaussian filter.

### Computation of mean heading vectors

We treated each heading measurement as a unit vector, and computed the mean of these heading vectors over a given window length of the fly walking (e.g. 60 s in Figure 1D). Heading measurements associated with time points in the analysis window where the fly was standing still (forward velocity < 0.5 mm/s) were omitted from contributing to the mean heading vector calculated in that window because when the fly stands still, its heading is constant for trivial reasons. We slid the analysis window over the heading time series within each trial (e.g. 60 s closed-loop bar in Figure 1), by 1 s increments, and calculated the mean-heading vector at each position, ultimately plotting the distribution of all mean heading vectors for a given trial type in a 2-dimensional polar histogram. Data near the periphery of such a histogram indicate walking along a consistent bearing whereas data near the middle indicate many changes in heading over the analyzed time window (23). We additionally calculated the mean vector length in each polar plot and then plotted the distribution of mean heading vector magnitudes of polar plots generated with analysis windows of different lengths (Figure S1B, Figure S6, Figure S11). Such a plot acts as a measure of how well flies maintain their heading over different timescales because if the mean vector of polar plots remains high with, for example, 400 s analysis windows, this indicates that flies must be maintaining consistent walking angles over 6+ minutes.

### Computing turning and forward walking as a function of distance to goal

Analyzing data from before and after bar jumps (−20 to 40 s), we computed 2D histograms of turning or forward walking velocity, as a function of distance to goal heading (Figures 2F, 2I, 4F, 4I, 5I and 5L). Like in all plots analyzing bar jump data, we operationally defined the goal heading as the mean heading 10 s before bar jumps. We also computed the mean turning signal as a function of distance to goal (black trace). One can plot these data with different delays between the turning velocity signal and the distance to goal signal. In all figures, we plot the data using the delay between the motor signal (e.g. turning or forward velocity) and the distance to goal signal at which the fly turned strongest as a function of distance to goal. This optimal delay was 350 ms for Figures 2, 4, and is consistent with the idea that the fly first processes information about the position of the bar, as well as other sensory and internal inputs, to compute its heading, and then translates this heading signal through further processing (for example, via the model in Figure 3) into a turning command. For Figure 5, we found that the turning response was strongest at 100 ms after the distance-to-goal E-PG phase signal, consistent with the E-PG phase signal (which is derived from the fly’s internal heading estimate) being closer to the fly’s motor command to turn than is a visual stimulus. We used 300 ms-Gaussian-filtered turning and forward walking signals for these analyses to smooth the turning and forward walking signals. The delays reported here are therefore expected to be approximate.

### Statistics

Statistical tests are indicated in the relevant figures. 95% confidence intervals were computed by bootstrapping.

---

**Supplemental Table 1.** Individually-labeled neurons from three E-PG Gal4 lines. Summary of the entire multicolor flip out data set. Each row represents an individual neuron. Information about the neuropil to which each neuron projects is shown in the PB (protocerebral bridge), EB (ellipsoid body) and Gall columns, respectively, as well as a fourth column (Other Neuropil) for other structures (when tracing neurons from the PB, EB or Gall, no innervation was found in other neuropil). We use a previously-published numbering scheme (9), which differs from others in the literature (31). Briefly, PB glomeruli are numbered 1-9 from left to right on each side. Glomeruli on the left side are preceded by ‘L’, and glomeruli on the right side are preceded by ‘R’. EB tiles are numbered 1-8, starting from the ventral-most tile going clockwise, when viewing the EB from the posterior side. EB wedges, two of which comprise one tile, are numbered according to their tile, as well as the half of the tile that they innervate: wedges innervating the counterclockwise-half of a given tile are labeled ‘L’, and wedges innervating the clockwise-half are labeled ‘R’. Using this numbering scheme, E-PG neurons innervate wedges and glomeruli with matching numbers and sides (e.g. an E-PG neuron innervating wedge 4L in the EB projects to glomerulus L4 in the PB) (31). In addition, some E-PG neurons innervate a demi-wedge in the EB (31); these are labeled as “demi” in the EB column, with “CW” and “CCW” indicating which half of the wedge is innervated. CW: clockwise. CCW: counterclockwise. D: dorsal. V: ventral. FLPG5 refers to the specific Flp recombinase used for low density neuron labeling (45). Note that the vast majority of (if not all) neurons identified by multicolor flip out, in all Gal4 lines, were consistent with the known anatomy of E-PGs.

| Gal4 Line | Cell | Brain | EB wedge | PB glomerulus | Gall | Other Neuropil | consistent with EPG | unambiguously EPG | comments |
| --- | --- | --- | --- | --- | --- | --- | --- | --- | --- |
| 27F02 | 1 | 1(FLPG5) | 3L-demi (CW-half) | L3 | D-right |  | y | y |  |
| 27F02 | 2 | 1(FLPG5) | 4L | L4 | V-right |  | y | y |  |
| 27F02 | 3 | 1(FLPG5) | 4R | R4 | ?-left |  | y | (y) |  |
| 27F02 | 4 | 1(FLPG5) | 5R | R5 | ?-left |  | y | (y) |  |
| 27F02 | 5 | 1(FLPG5) | 7L | L7 | D-right |  | y | y |  |
| 27F02 | 6 | 1(FLPG5) | 8L | L8 | V-right |  | y | y |  |
| 27F02 | 7 | 1(FLPG5) | 1L | L9 | D-right |  | y | y |  |
| 27F02 | 8 | 2(FLPG5) | 3L | L3 | D-right |  | y | y |  |
| 27F02 | 9 | 2(FLPG5) | 5L-demi (CW-half) | L5 | D-right |  | y | y |  |
| 27F02 | 10 | 2(FLPG5) | 7L | L7 | D-right |  | y | y |  |
| 27F02 | 11 | 2(FLPG5) | 1L | L9 | D-right |  | y | y |  |
| 27F02 | 12 | 2(FLPG5) | 5R | R5 | D-left |  | y | y |  |
| 27F02 | 13 | 2(FLPG5) | 8R | R8 | V-left |  | y | y |  |
| 27F02 | 14 | 3(FLPG5) | 4L | L4 | V-right |  | y | y |  |
| 27F02 | 15 | 3(FLPG5) | 7L-demi (CCW-half) | L7 | D-right |  | y | y |  |
| 27F02 | 16 | 3(FLPG5) | 8L | L8 | V-right |  | y | y |  |
| 27F02 | 17 | 3(FLPG5) | 2R | R2 | V-left |  | y | y |  |
| 27F02 | 18 | 3(FLPG5) | 7L-demi (CCW-half) | R7 | ?-left |  | y |  | EB wedge 7L connects to PB glomerulus 7R |
| 27F02 | 19 | 4(FLPG5) | 4L | L4 | V-right |  | y | y |  |
| 27F02 | 20 | 4(FLPG5) | 7L | L7 | D-right |  | y | y |  |
| 27F02 | 21 | 4(FLPG5) | 8R | R8 | V-left |  | y | y |  |
| 27F02 | 22 | 5(FLPG5) | 3L | L3 | D-right |  | y | y |  |
| **27F02** | 23 | 5(FLPG5) | 4R | R4 | V-left |  | y | y |  |
| **27F02** | 24 | 5(FLPG5) | 5R-demi<br>(CCW-half) | R5 | D-left |  | y | y | 1/4 to 1/2<br>volume of PB<br>glomerulus is<br>innervated |
| **27F02** | 25 | 5(FLPG5) | 6R | R6 | V-left |  | y | y |  |
| **27F02** | 26 | 5(FLPG5) | 7L | R7 | V-left |  | y |  | EB wedge 7L<br>connects to PB<br>glomerulus 7R |
| **total** | **26** |  |  |  |  |  | **26/26** | **24/26** |  |
| **77E05** | 1 | 1(FLPG5) | 2L | L2 | V-right |  | y | y |  |
| **77E05** | 2 | 1(FLPG5) | 5L-demi<br>(CCW-half) | L5 | D-right |  | y | y |  |
| **77E05** | 3 | 1(FLPG5) | 4R-demi<br>(CW-half) | R4 | V-left |  | y | y |  |
| **77E05** | 4 | 2(FLPG5) | 2L | L2 | V-right |  | y | y |  |
| **77E05** | 5 | 2(FLPG5) | 4R | R4 | V-left |  | y | y |  |
| **77E05** | 6 | 2(FLPG5) | 5R-demi<br>(CCW-half) | R5 | D-left |  | y | y |  |
| **77E05** | 7 | 3(FLPG5) | 4L-demi<br>(CCW-half) | L4 | V-right |  | y | y |  |
| **77E05** | 8 | 3(FLPG5) | 4L-demi<br>(CW-half) | L4 | V-right |  | y | y |  |
| **77E05** | 9 | 3(FLPG5) | 6L-demi<br>(CCW-half) | L6 | V-right |  | y | y |  |
| **77E05** | 10 | 3(FLPG5) | 2R | R2 | V-left |  | y | y |  |
| **77E05** | 11 | 3(FLPG5) | 4R-demi<br>(CCW-half) | R4 | V-left |  | y | y |  |
| **77E05** | 12 | 4(FLPG5) | 3L | L3 | D-right |  | y | y |  |
| **77E05** | 13 | 4(FLPG5) | 4L | L4 | V-right |  | y | y |  |
| **77E05** | 14 | 4(FLPG5) | 6R | R6 | V-left |  | y | y |  |
| **77E05** | 15 | 5(FLPG5) | 4L | L4 | V-right |  | y | y |  |
| **total** | **15** |  |  |  |  |  | **15/15** | **15/15** |  |
| **60D05** | 1 | 1(FLPG5) | 3L | L3 | D-right |  | y | y |  |
| **60D05** | 2 | 1(FLPG5) | 4L | L4 | V-right |  | y | y |  |
| **60D05** | 3 | 1(FLPG5) | 1L-demi<br>(CCW-half) | L9 | D-right |  | y | y |  |
| **60D05** | 4 | 1(FLPG5) | 2R-demi<br>(CW-half) | R2 | V-left |  | y | y |  |
| **60D05** | 5 | 1(FLPG5) | 8R-demi<br>(CW-half) | R8 | V-left |  | y | y |  |
| **60D05** | 6 | 2(FLPG5) | 3R | R3 | D-left |  | y | y |  |
| **60D05** | 7 | 2(FLPG5) | 5R | R5 | D-left |  | y | y |  |
| **60D05** | 8 | 2(FLPG5) | 7R | R7 | D-left |  | y | y |  |
| **60D05** | 9 | 3(FLPG5) | 5L-demi<br>(CW-half) | L5 | D-tight |  | y | y |  |
| **60D05** | 10 | 3(FLPG5) | 7R | R7 | D-left |  | y | y |  |
| **60D05** | 11 | 4(FLPG5) | 4L | L4 | V-right |  | y | y |  |
| **60D05** | 12 | 4(FLPG5) | 7L | L7 | D-right |  | y | y |  |
| **60D05** | 13 | 4(FLPG5) | 2R | R2 | V-right |  | y | y |  |
| **60D05** | 14 | 5(FLPG5) | 7L | L7 | D-right |  | y | y |  |
| **60D05** | 15 | 5(FLPG5) | 1L | L9 | D-right |  | y | y |  |
| **60D05** | 16 | 5(FLPG5) | 5R-demi<br>(CW-half) | R5 | D-left |  | y | y |  |
| **60D05** | 17 | 5(FLPG5) | 8R | R8 | V-left |  | y | y |  |
| **total** | **17** |  |  |  |  |  | **17/17** | **17/17** |  |

**Supplemental Table 2.** Genotypes and experimental parameters used in each figure. X-Gal4 represents three different transgenes: 60D05-Gal4, 77E05-Gal4, 27F02-Gal4. Trial orders were randomly permuted for each recording.

| Dataset | Genotype | Experiment | Figure # | # Flies | # Trials |
| --- | --- | --- | --- | --- | --- |
| 1 | Canton-S | 60 min closed-loop bar, 10 min dark | Fig. 1, S1 | 14 flies | NA |
| 2 | Canton-S | Closed-loop bar with discontinuous bar jumps every 3 min | Fig. 2, S2, S3 | 14 flies | 7 trials each of -90°, +90° and 180° rotations |
| 3 | Canton-S | 3 min closed-loop bar, 30 s dark, closed-loop bar reappears at random offset | Fig. 2C | 12 flies | 16 trials |
| 4 | + ; + ; 60D05-Gal4 / UAS-GCaMP6f | Closed-loop bar with discontinuous bar jumps every 2 min, with imaging | Fig. S4A-B | 22 flies | 2 trials each of -90°, +90°, 180° per recording (1 recording per fly, except 2 flies with 2 recordings, 1 fly with 3, and 1 fly with 4) |
| 5 | + ; + ; 60D05-Gal4 / UAS-GCaMP6f | 30 min continuous closed-loop bar, with imaging | Fig. S4C-D | 11 flies | NA |
| 6a | + / w ; 60D05-LexA / LexAOp-GCaMP6f ; X-Gal4 / UAS-*shibire*<sup>ts</sup> | 100 s each of continuous closed-loop bar and dark, with imaging | Fig. S5 | 10 flies (x3) | 1 trial at 26°C and 34°C |
| 6b | + / w ; 60D05-LexA / LexAOp-GCaMP6f ; X-Gal4 / + | 100 s each of continuous closed-loop bar and dark, with imaging | Fig. S5 | 10 flies (x3) | 1 trial at 26°C and 34°C |
| 6c | + / w ; 60D05-LexA / LexAOp-GCaMP6f ; + / UAS- *shibire*<sup>ts</sup> | 100 s each of continuous closed-loop bar and dark, with imaging | Fig. S5 | 10 flies | 1 trial at 26°C and 34°C |
| 7a | + / w ; 60D05-LexA / LexAOp-GCaMP6f ; X-Gal4 / UAS-*shibire*<sup>ts</sup> | 30 min closed-loop bar at 34°C | Fig. 4B-C, S6 | 10 flies (x3) | NA |
| 7b | + / w ; 60D05-LexA / LexAOp-GCaMP6f ; X-Gal4 / + | 30 min closed-loop bar at 34°C | Fig. 4B-C, S6 | 10 flies (x3) | NA |
| 7c | + / w ; 60D05-LexA / LexAOp-GCaMP6f ; + / UAS- *shibire*<sup>ts</sup> | 30 min closed-loop bar at 34°C | Fig. 4B-C, S6 | 10 flies | NA |
| 8a | + / w ; 60D05-LexA / LexAOp-GCaMP6f ; X-Gal4 / UAS-*shibire*<sup>ts</sup> | Closed-loop bar with discontinuous bar jumps | Fig. 4D-I, S7-9 | 10 flies | 14 trials each of -90°, +90° and 180° for half the |
|  |  | every 90 s |  | (x3) | flies, 14 trials each of -90° and +90° for other half of flies |
| 8b | + / w ; 60D05-LexA / LexAOp-GCaMP6f ; X-Gal4 / + | Closed-loop bar with discontinuous bar jumps every 90 s | Fig. 4D-I, S7-9 | 10 flies (x3) | 14 trials each of -90°, +90° and 180° for half the flies, 14 trials each of -90° and +90° for other half of flies |
| 8c | + / w ; 60D05-LexA / LexAOp-GCaMP6f ; + / UAS- *shibire<sup>ts</sup>* | Closed-loop bar with discontinuous bar jumps every 90 s | Fig. 4D-I, S7-9 | 10 flies | 14 trials each of -90°, +90° and 180° for half the flies, 14 trials each of -90° and +90° for other half of flies |
| 9 | 57C10-FLPG5 / + ; + ; 10xUAS-FRT.stop-myr::smGdP-HA, UAS-FRT.stop-myr::smGdP-V5-THS-10xUAS-FRT.stop-myr::smGdP-FLAG / X-Gal4 | Immunohistochemical labeling of individual neurons with different colors | Fig. S10, Supp. Table 1 | 5 flies |  |
| 10 | + ; + ; 60D05-Gal4 / UAS-GCaMP6f | 10 min closed-loop bar and 10 min dark, with imaging | Fig. S11 | 11 flies | NA |
| 11c | + / w ; 60D05-LexA / LexAOp-GCaMP6f ; VT032906-Gal4 / UAS-P2X2 | Dark, with ATP stimulations every ~120 s (manually triggered) , with imaging | Fig. 5, S12-13 | 17 flies | 11-16 stimulations per recording, 1 recording per fly |
| 11e | + / w ; 60D05-LexA / LexAOp-GCaMP6f ; + / UAS-P2X2 | Dark, with ATP stimulations every ~120 s (manually triggered) , with imaging | Fig. S12-13 | 15 flies | 14-15 stimulations per recording, 1 recording per fly |

**Supplemental Table 3.** Origin of each genotype used. BDSC: Bloomington Drosophila Stock Center, VDRC: Vienna Drosophila Resource Center.

| Transgene | Origin |
| --- | --- |
| Canton-S | Martin Heisenberg,<br>through Michael<br>Dickinson |
| pJFRC99-20XUAS-IVS-Syn21-Shibire-<br>ts1-p10 inserted at VK00005 | Gerald Rubin |
| UAS- P2X<sub>2</sub> | Vanessa Ruta |
| LexAOp-GCaMP6f | BDSC #44277 |
| UAS-GCaMP6f | BDSC #52869 |
| VT032906-Gal4 | VDRC #202537 |
| VT020739-Gal4 | VDRC #201501 |
| 60D05-LexA | BDSC #52867 |
| 60D05-Gal4 | BDSC #39247 |
| 77E05-Gal4 | BDSC #48338 |
| 27F02-Gal4 | BDSC #48072 |
| 57C10-FLPL / + ; + ; 10xUAS-FRT.stop-myr::smGdP-HA, UAS-FRT.stop-myr::smGdP-V5-THS-10xUAS-FRT.stop-myr::smGdP-FLAG | BDSC #64087 |
| 57C10-FLPG5 / + ; + ; 10xUAS-FRT.stop-myr::smGdP-HA, UAS-FRT.stop-myr::smGdP-V5-THS-10xUAS-FRT.stop-myr::smGdP-FLAG | BDSC #64088 |

**Movie S1. Example fly turning to bring its E-PG heading signal back to its position in the protocerebral bridge prior to neural stimulation.**

Movie of a fly walking in constant darkness (bottom). Two-photon imaging in the protocerebral bridge of E-PG neurons expressing GCaMP6f (top). The left bridge is shown on the right, and the right bridge on the left, since the fly is viewed from the front. We stimulated E-PGs via P-ENs (see *Methods*) by expressing in P-ENs the ATP-gated cation channel P2X2 and releasing ATP locally via a pipette (shown in red) on 1-2 glomeruli in the protocerebral bridge. Stimulating P-ENs reliably relocates the E-PG activity peaks approximately 1 glomerulus medial to the stimulated position (9), consistent with P-ENs functionally exciting E-PGs and their known projection anatomy (31). We added a red dot to the movie to appear at approximately the time when a pressure pulse releases ATP from the pipette. The red dot lasts 2 seconds in the movie, to ensure it is salient and visible to the viewer; the actual pressure pulse lasted only 20-50 ms. All other movie components aside from the red dot (e.g., brain activity and fly behavior) were aligned using timestamps from the behavioral camera and the two photon frame triggers in python 2.7.

**Movie S2. Same as Movie S1 but at 2x speed.**

